# Human Fis1 directly interacts with Drp1 in an evolutionarily conserved manner to promote mitochondrial fission

**DOI:** 10.1101/2023.05.03.539292

**Authors:** Kelsey A. Nolden, Megan C. Harwig, R. Blake Hill

**Affiliations:** Department of Biochemistry, Medical College of Wisconsin, Milwaukee, WI 53226 (USA)

**Keywords:** *mitochondria*, mitochondrial dynamics, fission, *protein-protein interaction*, *nuclear magnetic resonance (NMR)*, microscale thermophoresis (MST), autoinhibition, *biophysics*, *cell biology*, *confocal microscopy*

## Abstract

Mitochondrial Fission Protein 1 (Fis1) and Dynamin Related Protein 1 (Drp1) are the only two proteins evolutionarily conserved for mitochondrial fission, and directly interact in *S. cerevisiae* to facilitate membrane scission. However, it remains unclear if a direct interaction is conserved in higher eukaryotes as other Drp1 recruiters, not present in yeast, are known. Using NMR, differential scanning fluorimetry, and microscale thermophoresis, we determined that human Fis1 directly interacts with human Drp1 (*K_D_* = 12-68 µM), and appears to prevent Drp1 assembly, but not GTP hydrolysis. Similar to yeast, the Fis1-Drp1 interaction appears governed by two structural features of Fis1: its N-terminal arm and a conserved surface. Alanine scanning mutagenesis of the arm identified both loss- and gain-of-function alleles with mitochondrial morphologies ranging from highly elongated (N6A) to highly fragmented (E7A) demonstrating a profound ability of Fis1 to govern morphology in human cells. An integrated analysis identified a conserved Fis1 residue, Y76, that upon substitution to alanine, but not phenylalanine, also caused highly fragmented mitochondria. The similar phenotypic effects of the E7A and Y76A substitutions, along with NMR data, support that intramolecular interactions occur between the arm and a conserved surface on Fis1 to promote Drp1-mediated fission as in *S. cerevisiae*. These findings indicate that some aspects of Drp1-mediated fission in humans derive from direct Fis1-Drp1 interactions that are conserved across eukaryotes.

## Introduction

Proper mitochondrial and cellular function requires mitochondrial fission and fusion, the dynamic processes by which mitochondria split into separate daughter organelles, or merge into interconnected networks (1–4). Mitochondrial fission is performed by Dynamin Related Protein 1, a cytosolic large GTPase enzyme within the dynamin superfamily of proteins (5–7). Classically, this family of proteins is associated with modulating membrane fission and fusion events, targeting pathogens for removal via the innate immune response, and cytoskeletal organization. Unlike small GTPases which require GTPase-activating proteins (GAPs) or guanine exchange factors (GEFs), members of the dynamin superfamily of proteins do not require additional proteins to stimulate their activity. Instead, homotypic interactions typically driven by protein self-assembly stimulate nucleotide hydrolysis and subsequent activity (8, 9).

For Drp1, this involves its recruitment to the mitochondrion (or peroxisome) via tail-anchored proteins resident to the outer membrane, likely enriched at ER-mitochondrial or lysosomal-mitochondrial contact sites (10–12). Known Drp1 recruiting proteins in mammals include Fis1, Mff, MiD49, and MiD51 (13–19). Of these, only Fis1 and Drp1 are evolutionarily conserved across all eukaryotes. Like Drp1 (20, 21), *Fis1*^-/-^ mice are embryonic lethal, suggesting the protein performs critical cellular functions (22). In *Saccharomyces cerevisiae*, where Fis1 was first discovered, Fis1p directly interacts with the yeast proteins Mdv1p/Caf4p and the yeast Drp1 (Dnm1p) to recruit the GTPase to the outer mitochondrial membrane (23–28). In structural studies of yeast Fis1p, the first sixteen residues, called the Fis1 arm, physically occludes a concave surface created by its tetratricopeptide (TPR) domain (29, 30). This surface is lined with evolutionarily conserved residues suggesting the arm may act intramolecularly in an autoinhibitory manner to prevent access to a binding region. Consistent with this model, a direct yeast Fis1p-Dnm1p interaction is enhanced upon removal of the Fis1 arm (31). Given these proteins are conserved across species, it is reasonable to consider that a human Fis1 arm acts in a similar manner. However, the human Fis1 arm is eight residues shorter and in NMR structures(32), it is disordered and does not form intramolecular contacts with the conserved surface as in yeast structures (29, 30).

Despite this, we recently showed the human Fis1 arm can adopt an intramolecular conformation, similar to the yeast protein (33). Further, this conformation occludes well-conserved Fis1p residues that disrupt yeast Fis1p-Dnm1p binding when mutated. In mammalian cells, human Fis1 and Drp1 do appear to interact via FRET and coimmunoprecipitation, although the latter required crosslinking (15, 34, 35). Overexpression of human Fis1 stimulates Drp1 localization to mitochondria and increases mitochondrial fragmentation (16, 18, 33, 36), which is ablated upon co-expression of a dominant-negative Drp1 K38A variant (16, 18, 36). Recently, Fis1 was also shown to participate in Drp1-mediated fission events that selectively occurred at the periphery to remove damaged mitochondria. In all these studies, it is unknown if they were mediated by direct Fis1-Drp1 interactions (11), and many have questioned the role of Fis1 in Drp1-mediated mitochondrial fission. Genetic silencing of Fis1 does not elongate mitochondria in all cell types (35, 37) and human Drp1 can be recruited by other mitochondrial membrane-anchored proteins including Mff, MiD49, and MiD51, with Mff appearing to be the primary recruiter of Drp1 (1, 13, 37–42). Moreover, the GTPase activity of Drp1 is not affected by Fis1(36), and no known mammalian homologs of the essential yeast Fis1p-Dnm1p adaptor protein, Mdv1p, exist (28, 43). In yeast, loss of Fis1p can be functionally replaced by adding a mitochondrial tether to the cytoplasmic adaptor Mdv1p(42). Transfection of human Drp1 with Mff and Mids, but not Fis1, fragments mitochondrial in yeast lacking the native fission components(42). Also, human Fis1 inhibits the mitochondrial fusion machinery (46), is involved in endoplasmic reticulum-mediated apoptosis independent of fragmentation(47–49), and recruits the pro-mitophagic TBC1D15/17, a GTPase Rab7a activator (35, 50–53). Collectively, these findings suggest human Fis1 function has diverged considerably from yeast.

Given these discrepancies, we sought to determine whether Fis1 is involved in Drp1-dependent fission. We report that human Fis1 maintains a direct interaction with Drp1 via highly conserved Fis1 residues, and this interaction appears to be governed by the Fis1 arm as its removal revealed a latent interaction with Drp1 *in vitro*. Single amino acid substitutions to the Fis1 arm or conserved residues cause both gain- and loss-of-function behavior that indicates a dramatic tuning of mitochondrial morphology can be achieved with single amino acid substitutions to Fis1. The analysis of these data shows that Fis1 can support Drp1-mediated mitochondrial fission in an evolutionarily conserved manner.

## Results

### Human Fis1 interacts with its own N-terminal region like yeast Fis1p

Given the ability of the human Fis1 arm (residues 1-8) to act intramolecularly (33), we asked whether an exogenously added synthetic peptide comprised of the eight-residue arm could bind to a recombinant version of Fis1 lacking the arm. For this, increasing amounts of a synthetic Fis1 arm peptide (M_1_EAVLNEL_8_) were titrated into an armless human Fis1 construct (Fis1^9-125^ or Fis1ΔN) uniformly labeled with ^15^N and followed by heteronuclear single quantum coherence (HSQC) NMR experiments. Overlays of the resulting ^1^H-^15^N HSQC spectra revealed chemical shift perturbations of only a subset of residues consistent with specific binding of the peptide. (**Figure 1A,B**). Plotting the first principal component from TREND analysis clearly showed saturation was not achieved even at 60-fold excess of peptide (**Figure 1C**). A global fit to the data gave an apparent *K_D_* of at least 5 mM but was not well determined. Regardless, the differences in chemical shifts (Δδ) between Fis1ΔN alone and at the highest concentration of peptide (3.5 mM) indicated that the greatest perturbations upon binding were from residues A107, L110, R83, L77, L84, R112, Y76, A78, and V79 (**Equation 3**, **Figure 1D**). Displaying all chemical shift perturbations onto the NMR structure of Fis1 (PDB: 1PC2 truncated to residues 9-125; **Figure 1E**) showed the most perturbed residues clustered on the concave surface of Fis1, a highly conserved region as determined by ConSurf analysis (54) (**Figure S1**), and strikingly similar to the surface where the Fis1p arm binds in the yeast molecule (29, 30).

**Figure 1.**
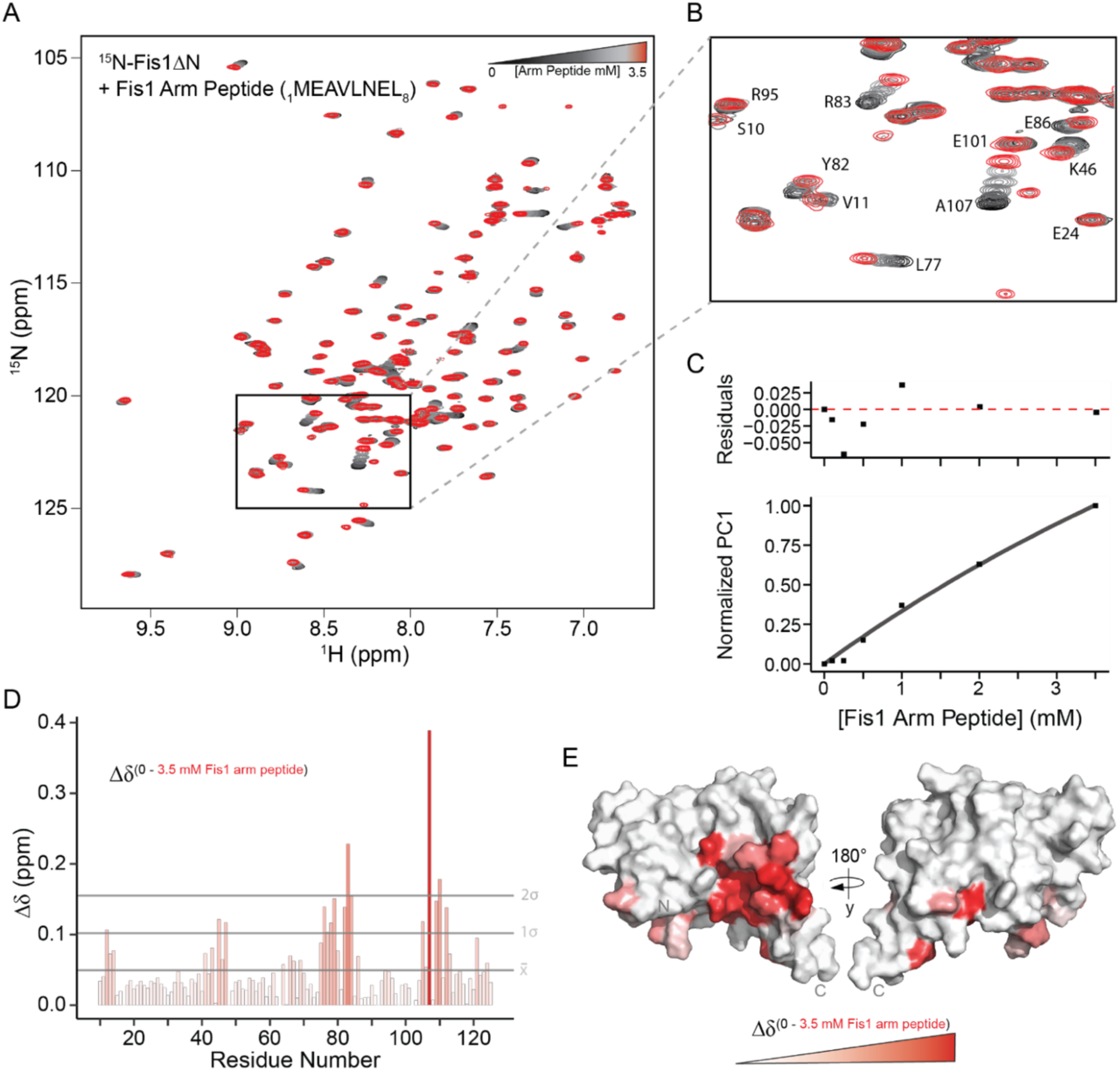
Fis1ΔN can bind a synthetic peptide version of the Fis1 N-terminal arm. *A.* ^1^H, ^15^N HSQC spectral overlays of 50 µM ^15^N-Fis1^9-125^ (ΔN) in the presence of increasing Fis1 N-terminal arm peptide (0-3 mM; 0, 100, 250, 500, 2000, 3500 µM) in a black-to-grey-to-red coloring scheme. *B.* Enlarged region of HSQC spectral overlays from (*A*) *C.* Affinity determination of arm peptide binding to Fis1ΔN using TREND analysis on all spectra from (*A*) and fitting the resulting PC1 values as a function of peptide concentration indicating an app. *K_D_* > 5 mM. *D.* Chemical shift perturbations of Fis1ΔN alone or in the presence of 3.5 mM N-terminal arm peptide (Δδ) are shown for each Fis1ΔN residue in a gradient fashion, where a redder color indicates a greater Δδ. The mean Δδ and one and two SD from the mean are indicated by horizontal lines. *E.* Fis1ΔN Δδ values displayed on a surface representation of the structure of Fis1 with the N-terminal arm removed (PDB: 1PC2^9-125^) in a gradient fashion that replicates the color scheme from (*D*).

### Removal of the Fis1 N-terminal arm reveals Drp1 binding in vitro

In yeast, the Fis1p arm inhibits Dnm1p binding, necessitating its removal for the detection of a robust Fis1p-Dnm1 interaction *in vitro* (31). Given human Fis1 can also bind its own N-terminal arm, we asked if the human Fis1 arm may similarly occlude Drp1 binding. To evaluate this possibility, we used differential scanning fluorimetry to monitor the intrinsic fluorescence of 30 µM Drp1 with increasing amounts of Fis1ΔN. In the absence of Drp1, the T_m_ of Fis1ΔN was found to be 77.5 ± 0.1 °C (**Figures 2A,S2C**), consistent with prior results (33). In the absence of Fis1ΔN, Drp1 thermally unfolds in two transitions with midpoints at 48.7 ± 0.2 °C and 76.8 ± 0.2 °C corresponding to the GTPase and stalk domains, respectively, as described previously (56, 57). Upon addition of increasing Fis1ΔN, only the second transition was altered and decreased in a concentration-dependent manner (**Figures 2A,S2C**). At equimolar concentrations (30 µM) of Fis1ΔN and Drp1, a single T_m_ for this transition was observed at 72.4 +/- 0.1 °C, a 5.1 °C decrease from the Fis1ΔN T_m_ and a 4.4 °C decrease from the Drp1 T_m_, suggesting the formation of a complex that likely alters the thermal unfolding profiles of both proteins (**Figures 2A,S2C,D**). In addition, this decrease in T_m_ appeared saturable as the concentration of Fis1ΔN increased (**Figure 2A**). These thermal melt data suggested a direct Fis1-Drp1 interaction.

**Figure 2.**
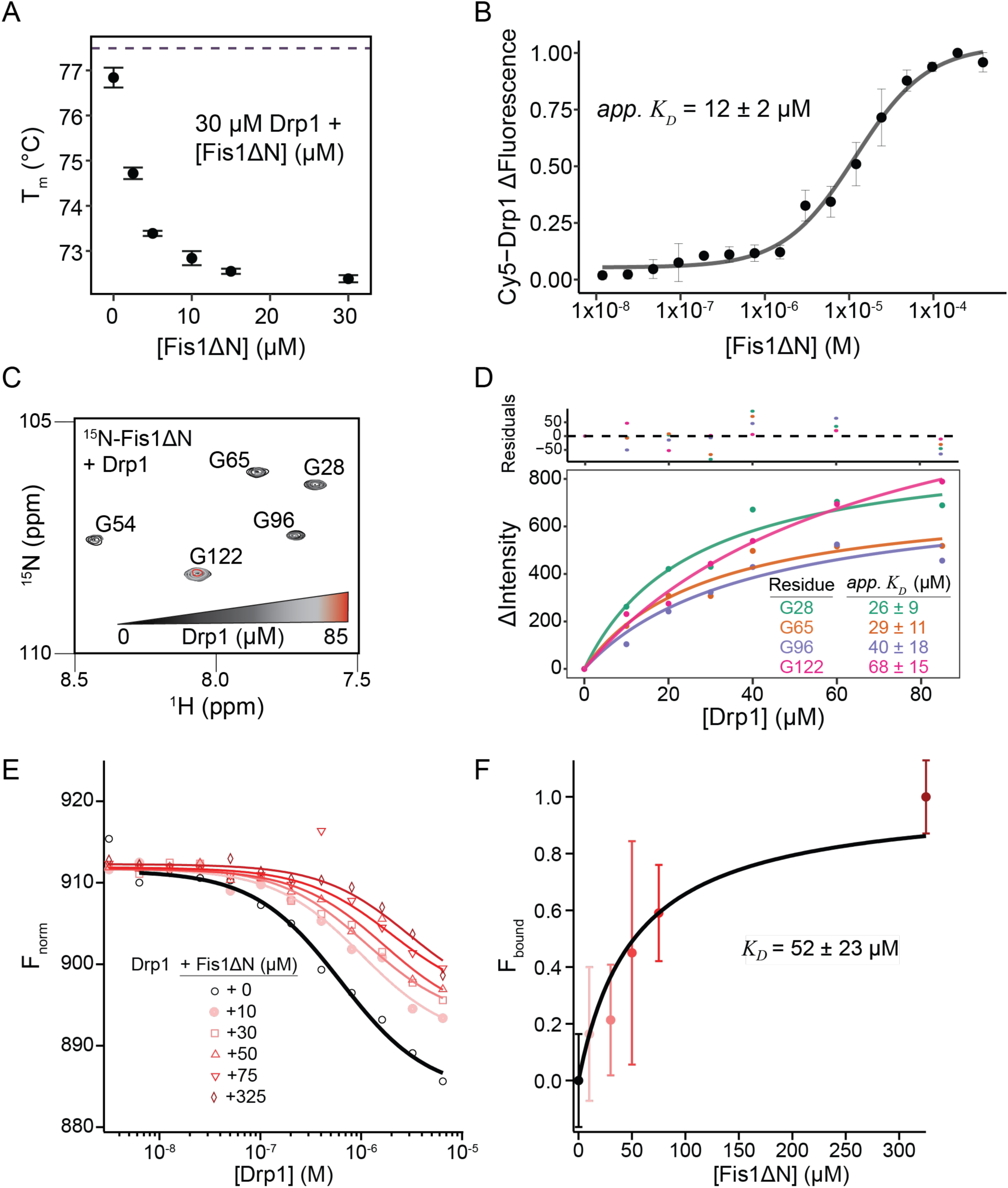
Removal of the Fis1 N-terminal arm reveals Drp1 binding and prevents Drp1 assembly *in vitro*. *A.* Box plot depicting the global unfolding temperature (T_m_) of 30 µM Drp1 in the presence of increasing Fis1ΔN (0-30 µM). T_m_ values determined as the temperature corresponding to the first derivative of the maximum fluorescence value. N=2 for each titration point. *B.* ΔFluorescence values of 80 nM Cy5-Drp1 in the presence of increasing Fis1ΔN as determined by microscale thermophoresis and fit to a single-site binding model to determine an apparent *K_D_* value. ΔFluorescence values normalized to a 0-1 scale to allow for comparisons and averaging between multiple experiments (n=3), error bars=SD. *C.* ^1^H, ^15^N HSQC spectral overlays of 50 µM ^15^N-Fis1ΔN^9-125^ in the presence of increasing Drp1 (0-85 µM) in a black-to-grey-to-red coloring scheme, full spectra in Figure S2E *D.* ΔIntensity values of select ^15^N-Fis1ΔN residues from (*C*) as a function of Drp1 concentration, fit to a single-site binding model to determine apparent binding affinities. *E.* Normalized fluorescence values (F_norm_) of 80 nM Cy5-Drp1 titrated with unlabeled Drp1 (0-103 µM) as a function of increasing concentrations of Fis1ΔN (0-325 µM) shown in a gradient coloring scheme from black (0 µM Fis1ΔN) to pink to red (325 µM Fis1ΔN). Data fit to an isodesmic model to determine effects of Fis1ΔN on Drp1 self-assembly. *F.* F_norm_ values from (E) converted to fraction bound (F_bound_) and plotted as a function of increasing FisΔN concentrations. Data fit to a one-site binding model to determine an apparent *K_D_* of Drp1 for Fis1ΔN. Error bars = SD of fit.

To test this further, Drp1 was fluorescently labeled with Cy5 for microscale thermophoresis (MST) experiments that utilize the change in thermophoretic behavior to detect binding (58, 59). In this experiment, unlabeled Fis1ΔN was titrated into Cy5-labeled Drp1 and the normalized changes in fluorescence intensity were monitored as a function of increasing Fis1ΔN. The resulting data were fit to a dose-response model to give an apparent binding affinity of *K_D_* = 12 ± 2 µM (**Figure 2B**). The inverse experiment in which unlabeled Drp1 was titrated into Cy5-Fis1ΔN compared favorably with an apparent binding affinity of 15 µM ± 5 µM (**Figure S2A**). We conclude that removal of the Fis1 arm allowed for ready identification of a direct interaction with Drp1, akin to the yeast system.

To potentially identify Fis1 residues mediating Drp1 binding, we titrated increasing amounts of Drp1 (0-85 µM) into 50 µM ^15^N-Fis1ΔN and monitored chemical shifts by 2D HSQC NMR as above (**Figures 2C,S2E**). As Drp1 concentrations increased, ^15^N-Fis1ΔN crosspeak intensity decreased, consistent with a strong interaction with Drp1, an 80 kDa protein that self-assembles into multimers and would be expected to decrease Fis1 signal upon interaction due to resonance broadening from the significant enhancement of the spin-spin relaxation rate. Plotting this intensity decrease for well-resolved glycine residues and fitting to a 1:1 binding model (**Equation 4**) gave apparent *K_D_* values ranging from 26-68 µM (**Figure 2D**). To assess the possibility that Drp1 binding may reflect non-specific binding due to exposure of Fis1’s TPR core, we tested whether ^15^N-Fis1ΔN can bind to BSA, a known promiscuous binder, and found no significant chemical shift perturbation or loss of intensity, suggesting no interaction (**Figure S3A**). Fis1ΔN was also found to interact with both Drp1 isoform 1 and 5 by 2D NMR, suggesting the interaction is likely not restricted to a single isoform (**Figure S3B**).

Drp1 self-assembly, as an amphitropic dynamin family member (60), is critical to its hydrolytic activity and mitochondrial fission. To determine if Fis1ΔN binding influenced Drp1 hydrolysis, we measured phosphate release upon incubation of Drp1 with GTP with Fis1ΔN at 50-fold excess and found no significant different in hydrolysis (**Figure S4**). This result was consistent across multiple preparations of protein and prior work (15). To determine Fis1ΔN influence on Drp1 assembly, we devised a microscale thermophoresis assay for Drp1 assembly with Cy5-Drp1 and monitored normalized fluorescence intensity upon titration with increasing amounts of unlabeled Drp1. The data are well fit to an isodesmic model for assembly with an apparent *K_D_* = 0.6 ± 0.1 µM. We then repeated this experiment in the presence of increasing concentrations of Fis1ΔN to determine if Fis1ΔN alters Drp1’s propensity to self-assemble. As Fis1ΔN concentration increased, the Drp1 self-assembly weakened (**Figure 2E**) in a concentration dependent manner. These data were fit to give apparent *K_D_* values for Drp1 assembly at each Fis1ΔN concentration with a nearly ten-fold decrease in apparent *K_D_* for Drp1 assembly at the highest Fis1 concentration (app *K_D_* = 3 ± 0.4 µM). We also fit these data to estimate an apparent *K_D_* for the Fis1ΔN-Drp1 interaction. For this, we converted the data in Figure 2E to fraction bound assuming 100% occupancy at the highest Fis1ΔN concentration of 325 µM. Fitting these data to a binding isotherm gave an apparent *K_D_* of 52 ± 23 µM (**Figure 2F**), strikingly consistent with the three other methods used to evaluate the Fis1ΔN-Drp1 interaction. These data suggest that Fis1ΔN decreases Drp1 self-assembly in solution without influencing hydrolysis.

### Fis1 arm residues influence mitochondrial morphology and Drp1 localization

Previously, we showed that Fis1 overexpression drives mitochondrial fission and clumping with increased recruitment of Drp1, all of which are significantly reduced with Fis1ΔN expression (33). Thus, removal of the Fis1 arm enhances Drp1 binding *in vitro*, but reduces Drp1 localization *in vivo*. To better understand these differences, we used an alanine scanning approach in which each of the eight residues within the Fis1 arm was sequentially replaced with an alanine (or glycine for A3). These eight variants were individually co-transfected with mitoYFP into human retinal pigmented epithelial (RPE) cells lacking the *Fis1* gene and mitochondrial morphology visualized by confocal microscopy (33) **(Figures 3A,S5**). For controls, wild type and Fis1 null cells were transfected with pcDNA alone. To quantify mitochondrial morphology, we used MitoGraph as previously described (33, 61, 62). Briefly, MitoGraph utilizes 3D confocal microscopy images to generate three-dimensional surface map models of mitochondrial networks within cells. In addition, MitoGraph computes a variety of metrics using graph theory, based on distances between mitochondrial ends and branch points, which can then be incorporated into a single MitoGraph Connectivity Score (MCS). This score reflects how fused or fragmented the networks are, with a higher value indicating a more elongated or interconnected network, and a lower value indicating a more fragmented network.

**Figure 3.**
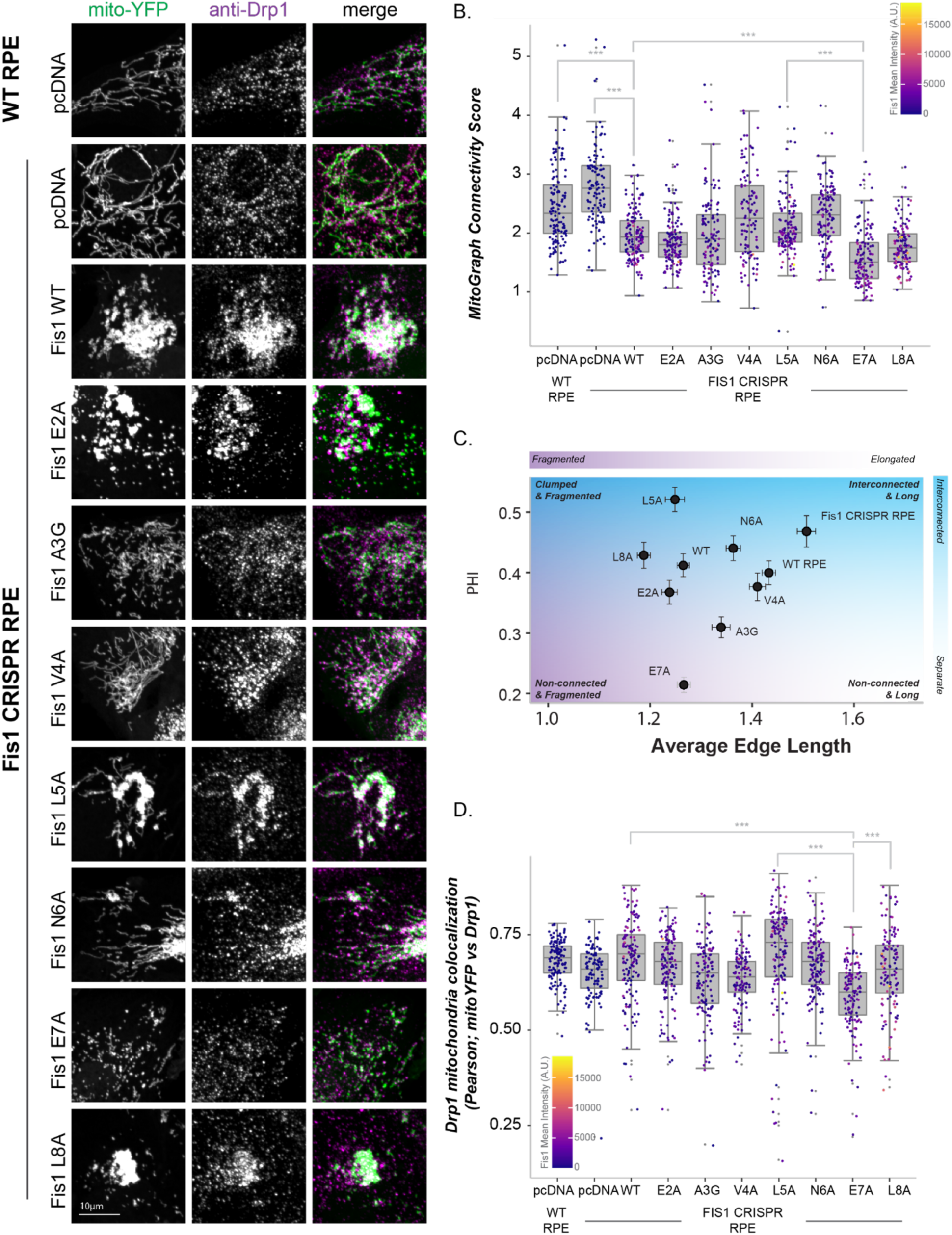
Alanine scanning of Fis1 arm alters mitochondrial morphology and Drp1 localization. *A-C*. WT or Fis1 KO retinal pigment epithelial (RPE) cells transfected with mitoYFP and either pcDNA, pcDNA-Fis1, or pcDNA-Fis1 point substitutions. *A.* Representative single-cell confocal images of anti-Drp1 and mitoYFP. Merged images show Drp1 (magenta) localization to the mitochondria (mitoYFP, green). *B*. Single cell z-stack images of mitoYFP transfected cells were segmented using MitoGraph on the mitoYFP channel and the MitoGraph Connectivity Score was calculated. Higher score is more elongated mitochondria. *C.* PHI vs. Average Edge Length scores calculated during MitoGraph analysis plotted with background pseudo-colored using a blue to purple gradient to represent four possible categories of mitochondrial network morphologies. Error bars = SEM. *D.* Colocalization between mitoYFP and anti-Drp1 from the same single cell z-stack images as in (*B*) measured using Pearson’s Correlation R-value. Data in (*B*) and (*D*) represented as boxplots with overlaid data points corresponding to the individual cells used for analysis. Gradient coloring in (*B*) and (*D*) represents Fis1 mean intensity by immunofluorescence of individual cells.

Consistent with prior results, Fis1 deletion resulted in significantly more elongated mitochondria (mean MCS increased from 2.4 to 2.7), whereas Fis1 overexpression resulted in more fragmented and collapsed networks (mean MCS decreased from 2.7 to 2.0) (**Figure 3A,B**). Dramatic changes to mitochondrial morphology were caused by single amino acid substitutions in the Fis1 arm that ranged from extremely fragmented mitochondria without clumping (E7A) or highly clumped organelles (L8A). These differences were quantified by MitoGraph (**Figures 3B, S6**) with the clumped morphologies being well captured by plotting the individual MitoGraph metrics Average Edge Length vs Phi (**Figure 3C**). The average edge length is the mean value of all individual mitochondrion’s length (called a connected component), and PHI is a measure of the size of the largest connected component to relative to the total mitochondrial network. This analysis revealed three main types of morphology: non-connected and fragmented (E7A), clumped and fragmented (wild-type, E2A, L5A, and L8A), and interconnected and long (WT RPE, Fis1 CRISPR RPE). The A3G, V4A, and N6A variants clustered to a fourth group representative of intermediate network morphologies. Overall, the profound changes to mitochondrial morphology did not arise from differences in Fis1 expression levels as variants expressed to same levels as wild type (**Figures 3B,S7**). Thus, small changes to Fis1 sequence have large changes on mitochondrial morphology.

To better understand the basis for Fis1 variants, Drp1 localization was measured using immunohistochemistry against Drp1 and quantified using a Pearson’s correlation coefficient with the mitoYFP signal (**Figure 3D**). In general, Ala substitutions that increased mitochondrial clumping (WT, E2A, L5A, and L8A) correlated well with Drp1 colocalization with L5A expression driving the largest increase of Drp1 localization to mitochondria. By contrast, Ala substitutions that decreased mitochondrial clumping (A3G and E7A) correlated poorly with Drp1 colocalization with E7A expression driving the largest decrease of Drp1 to mitochondria. Drp1 localization was well correlated with the MitoGraph PHI score (R^2^ =0.74, **Figure S8)** consistent with expression of ΔN (33) and mitochondrial clumping being both Fis1 and Drp1 dependent.

### E7 of Fis1 appears to interact with conserved residues but does not alter Drp1 affinity in vitro

Given the profound mitochondrial fragmentation upon expression of Fis1-E7A, we determined its impact on Fis1 conformation and interactions with Drp1. An overlay of ^1^H-^15^N HSQC spectra from uniformly labeled E7A and WT show only eight residues with significant chemical shift perturbations (L8, C41, V43, S45, K108, S10, W40, N48, Y76, **Figure 4A,B**) that occur in four regions of primary structure and cluster on the concave surface of Fis1 (**Figure 4C,D**). By MST, Fis1 E7A has essentially identical affinity for Drp1 as Fis1ΔN with a *K_D_* of 7 ± 1 µM (**Figure 4E**). Similarly, ^15^N-Fis1 E7A appears to bind Drp1 by 2D NMR (**Figure S9**). We infer from these data that the arm does not directly mediate interactions with Drp1, but E7A likely favors a conformation in which the arm is displaced from the concave surface facilitating Drp1 binding.

**Figure 4.**
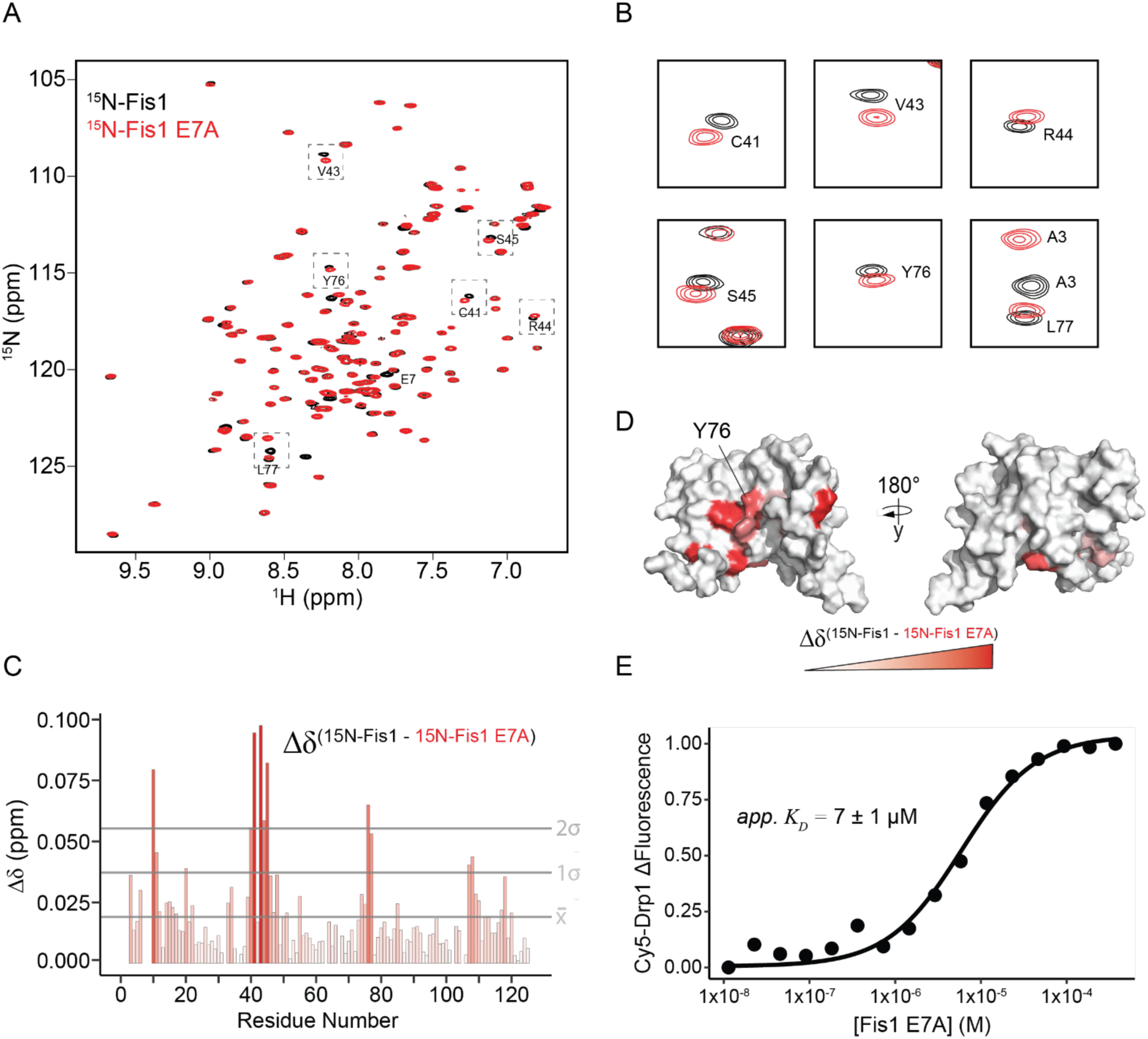
The Fis1 E7A substitution perturbs conserved Fis1 residues without disrupting Drp1 binding. *A.* ^1^H, ^15^N HSQC spectral overlays of 50 µM ^15^N-Fis1 and 50 µM ^15^N-Fis1 E7A with a subset of perturbed residues highlighted and enlarged in (*B*). *C.* Chemical shift perturbations (Δδ) between Fis1 and Fis1 E7A are shown for each Fis1 residue in a gradient fashion, where a redder color indicates a greater Δδ. Residue L8 omitted due to large perturbation (expected from substitution of neighboring residue). The mean Δδ and one and two SD from the mean are indicated by horizontal lines. *D.* Δδ values displayed on a surface representation of the structure of Fis1 (PDB: 1PC2^1-125^) in a gradient fashion that replicates the color scheme from (*C*). *E.* Normalized ΔFluorescence of Cy5-Drp1 in the presence of increasing unlabeled Fis1 E7A fit to a single-site binding model indicating an apparent *K_D_* = 7 ± 1 μM. N=1.

### Fis1-Y76A phenocopies Fis1-E7A in mitochondrial morphology and Drp1 localization

The concave surface of human Fis1 is lined with conserved residues (**Figure S1**, **Table S2**), which in yeast mediate pulldown interactions with Dnm1p (31). Integrating these two classes of residues (highly conserved and yeast-inspired) with residues with the greatest NMR signal loss upon Drp1 addition (**Figure S2E**) identified a handful of residues from the intersectionality of these classes (**Figure 5A**). This intersectionality is defined by four regions, three of which are shared by two classes and one shared by all three classes. A handful of amino acids fall into three of these regions with Y76 common to all classes. Constructs encoding these variants were individually transiently co-transfected with mitoYFP into human RPE Fis1*^-/-^* cells (**Figures 5B, S10**) and both mitochondrial morphology and Drp1-mitoYFP colocalization were evaluated as described above.

**Figure 5.**
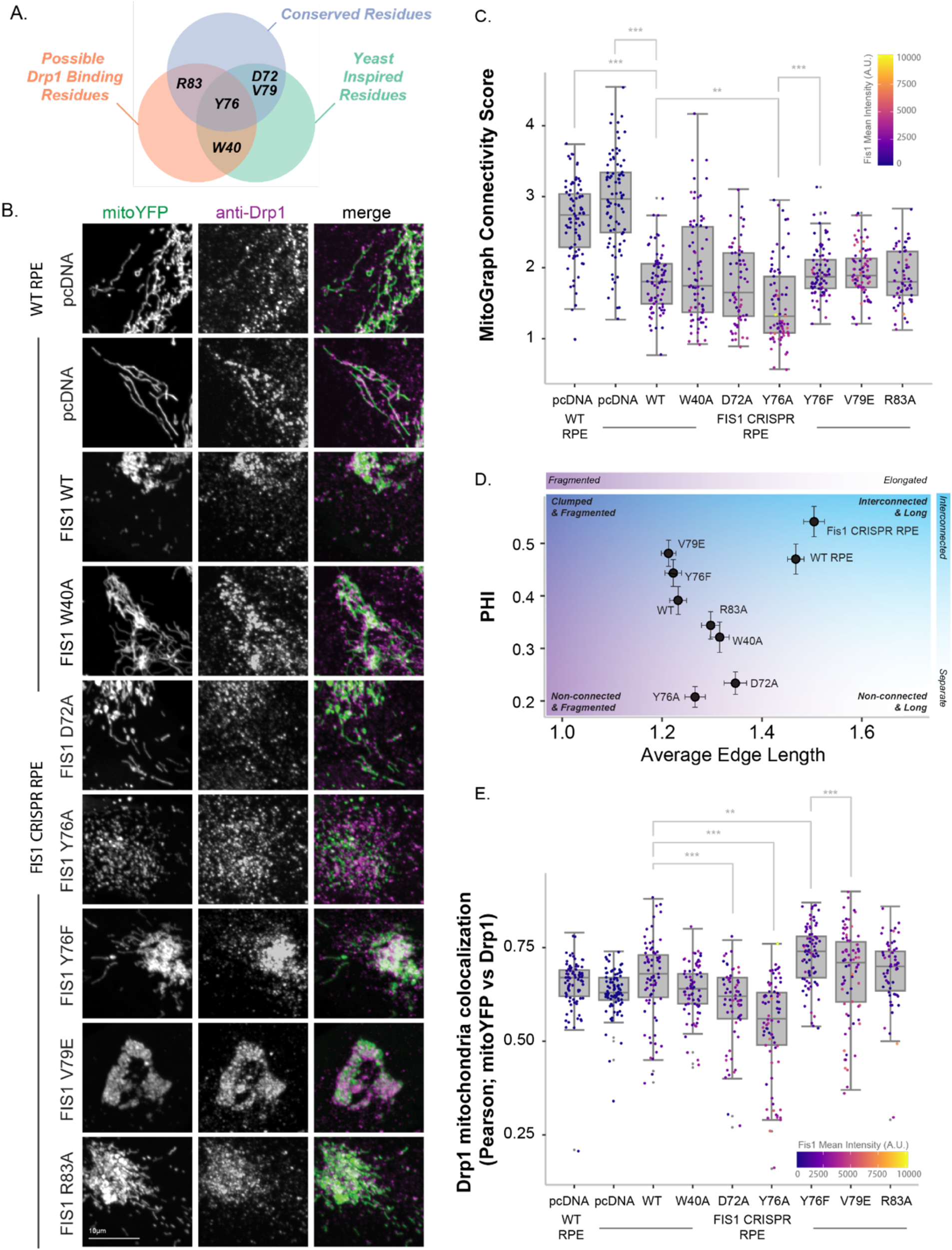
Substitution to conserved Fis1 residues alters mitochondrial fission and Drp1 localization. *A.* Venn Diagram identifying human Fis1 candidate residues for mutation that derive from comparing conserved non-TPR Fis1 residues, possible Drp1 binding residues (from Figure 2 analysis), and the orthologous human residue that induce phenotypes in yeast upon mutation. *B-D.* WT or Fis1 KO RPE cells transfected with mitoYFP and either pcDNA, pcDNA-Fis1, or pcDNA-Fis1 point substitutions. *B.* Representative single-cell confocal images of anti-Drp1 and mitoYFP. Merged images show Drp1 (magenta) localization to the mitochondria (mitoYFP; green). *C.* Single cell z-stack images of mitoYFP transfected cells were segmented using MitoGraph on the mitoYFP channel and the MitoGraph Connectivity Score was calculated. *D.* PHI vs. Average Edge Length scores calculated during MitoGraph analysis plotted with background pseudo-colored using a blue to purple gradient to represent four possible categories of mitochondrial network morphologies. Error bars = SEM. *E.* Colocalization between mitoYFP and anti-Drp1 from the same single cell z-stack images as in (*C*) measured using Pearson’s Correlation R-value. Data in (*C*) and (*E*) represented as boxplots with overlaid data points corresponding to the individual cells used for analysis. Gradient coloring in (*C*) and (*E*) represents Fis1 mean intensity by immunofluorescence of individual cells.

Human Fis1 variants whose orthologous position in yeast disrupted morphology (W40A, D72A) or Dnm1p interactions (Y76A, V79E, R83E) altered mitochondrial morphology to varying degrees (**Figure 5B-D,S10-11**). Notably, Fis1-Y76A was strikingly similar to E7A with extremely fragmented mitochondria without clumping. This similarity is shown in the Average Edge Length vs PHI correlation plot (**Figure 5D**), where Y76A shared nearly identical values with E7A in the non-connected and fragmented region. By contrast, Y76F in which the aromaticity of the side chain was preserved, clumped mitochondria more than wild-type Fis1 as captured by the PHI score, as did V79E. W40A, D72A, and R83A had intermediate phenotypes characterized by slightly elongated mitochondria with varying degrees of perinuclear clumping (**Figure 5D**). These changes did not arise from differences in Fis1 expression levels (**Figure 5C,S12**). Y76A expression also decreased Drp1 localization to mitochondria to a similar extent as E7A, whereas Y76F did not (**Figure 5E**). Thus, Y76A phenocopied E7A with both variants showing extreme mitochondrial fragmentation with less Drp1 colocalization. We interpret these data to suggest reciprocal interactions between the arm and concave surface are important for Drp1 recruitment and fission.

## Discussion

Although Fis1 is considered a fundamental component of the mitochondrial fission machinery in yeast, its role in human Drp1-mediated fission has remained uncertain. Consistent with prior results, transient overexpression of wild-type Fis1 drives mitochondrial fragmentation and perinuclear clumping (16, 18, 33, 35, 63). Using multiple biophysical techniques, we show a direct Fis1-Drp1 interaction. Our data support that two critical structural features of Fis1, the N-terminal arm and the TPR concave surface, are involved in this interaction, which appears to prevent Drp1 assembly. Titrations with an arm-derived peptide into Fis1ΔN show perturbations by NMR chemical shifts to residues in the TPR concave surface, suggesting that the arm interacts with this surface including residues Y76, R83, A107, AND L110 (**Figure 1**). Arm-TPR concave surface interactions are also present given that arm variant E7A shows strong chemical shift perturbations to similar concave surface residues, including Y76 and A107 (**Figure 4**). These data, along with prior NMR data upon arm deletion (33), are consistent with the Fis1 arm physically occluding access to the concave surface despite it being disordered in NMR structures(32). Thus, by occluding the concave surface, the arm acts intramolecularly to autoinhibit interactions with Drp1. This appears to be a kinetic effect as E7A does not significantly alter affinity with Drp1. Supporting this idea, Fis1 has a similar affinity for Drp1 (13 ± 3 µM) to Fis1ΔN by MST (**Figure S2B**) but only if a sufficient incubation time is used.

The consequences of these regions for Fis1 activity are demonstrated by profoundly different effects on mitochondrial morphology upon expression of single alanine substitutions. We observe two distinct changes to mitochondrial network morphologies—perinuclearly clumped or highly-fragmented and well-dispersed. This contrast is evident in comparing the E7A vs. L8A (**Figure 3**) and the Y76A vs. Y76F substitutions (**Figure 5**): L8A and Y76F cause mitochondrial perinuclear clumping similar to wild-type expression, whereas E7A and Y76A cause mitochondrial fragmentation without clumping. It may be that the arm directly binds to Drp1 to support Fis1-Drp1 interactions. However, wild-type, E7A, and ΔN all bind to Drp1 with similar affinities (**Figures 2,4,S2**). Previously we showed the Fis1 arm is in a dynamic equilibrium between at least two conformations, one that favors intramolecular interactions with the TPR concave surface and would be expected to prevent Drp1 binding and another conformation in which the arm is disordered and fully solvent exposed. We propose the variants studied here likely modulate this equilibrium, which is supported by changes in NMR chemicals shifts for E7A (**Figure 4**).

Fis1 variants also show profoundly different effects on mitochondrial localization of Drp1, which is increased for highly clumped, and decreased for highly fragmented, morphologies. Insight into these differences may be resolved from considerations of the Drp1 hydrolytic cycle in which GTP hydrolysis can catalyze membrane constriction, disassembly, and release from the membrane(64, 65). Thus, one plausible model is that hyper-fragmentation derives from a Fis1 conformation conducive to more efficient cycling of Drp1 on/off the mitochondria, which is favored by E7A/Y76A, but not L8A/Y76F, possibly by displacing the Fis1 arm to support productive Drp1 interactions. Productive Drp1 assembly occurs from the dimeric species (66) and it is tempting to speculate that Fis1 favors this species, sequestering dimeric Drp1 for productive assembly without futile cycling of GTP. This model would be consistent with our data including Fis1 influence on Drp1 self-assembly (**Figure 2E**), as well as Fis1ΔN’s ability to interact with the pathological obligate-dimer Drp1 G401S variant (**Figure S13**). However, this does not explain how wild-type and some variants (L5A, L8A, Y76F, and V79E) cause a clumped morphology. In our prior work in the yeast system, we identified a Fis1 variant, E78A, that caused uniform mitochondrial localization of Dnm1p from discrete punctate structures. Based on this work, we proposed a lattice-like model for assembly of the mitochondrial fission machinery in which the proper balance of interactions is necessary for fission. We speculate a lattice-like model could also explain our results here: the clumped morphology derives from an improper balance of interactions that prevents Drp1 disassembly resulting in clumping, and the hyper-fragmented morphology derives from interactions that facilitate more productive Drp1 lattice formation and disassembly to enhance fission.

If the Fis1 arm acts in an autoinhibitory manner to prevent Drp1 binding as our data suggests, it raises the question of how this inhibition may be relieved. Fis1 undergoes numerous post-translational modifications (67), of which phosphorylation by DNA-PKcs and Met enhance mitochondrial fission (68, 69), whereas ubiquitination (70, 71), SUMOylation (72), and acetylation (73, 74) decrease mitochondrial fission. While our data do not identify the mechanism of autoinhibition relief, significant chemical shift perturbations from the E7A substitution are localized to a conserved, non-canonical insert in the first TPR of Fis1 that comprises residues S45-K46-Y47 (the SKY insert) that have been recently identified as being important in Fis1 activity (75). Collectively, our data would support a model in which post-translational modifications to either arm or SKY residues could relieve autoinhibition. Another possibility is that Fis1 may require oligomerization as a necessary step in Drp1 recruitment (76). Consistent with this, previous work suggests that Fis1 α-helix_1_ (residues 11-32) is a negative regulator of Fis1 oligomerization (77). When this helix and the Fis1 arm is removed in human cells (Fis1^32-152^) it results in swollen punctate mitochondria and increased Drp1 localization that appears derived from increased Fis1 oligomerization. Similarly, removal of the N-terminal arm from yeast Fis1p also induces protein dimerization(78). These findings are consistent with our results here in which removal of a shorter stretch of human Fis1 (residues 1-8) increases Drp1 interactions *in vitro*, although how arm deletion influences human Fis1 oligomerization in cells is unknown. Fis1 oligomerization also appears to modulate MiD49 oligomerization as a necessary precursor for MiD49-mediated Drp1 recruitment, suggesting these two processes are linked (79). Fis1’s role in mitochondrial fission may also be through inhibition of the fusion machinery (46). Although our results here suggest Fis1 is capable of directly binding to Drp1, we cannot exclude that well-dispersed and highly fragmented mitochondria from Fis1 E7A and Fis1 Y76A may arise from enhanced interactions with the fusion proteins, especially given MFN2’s role in perinuclear clumping of mitochondria (80).

Notably, Fis1 variants with well-dispersed and fragmented mitochondria also had decreased Drp1, as determined by total Drp1 fluorescence intensity (**Figures S6,S11**), suggesting increased protein degradation. This could occur through increased rates of mitophagy, especially given Fis1’s proposed role in this process via its interactions with Drp1, TBC1D15, and STX17 (11, 35, 52, 53, 76, 79, 81). Indeed, a pivotal study found that mitochondrial fission facilitated the separation and removal of damaged mitochondria from the larger network by mitophagy, suggesting the processes of fission and mitophagy can occur sequentially (82). Emerging evidence also suggests that Fis1 may be the primary Drp1 recruiter under stress-induced fission events, likely as a precursor to mitophagy, whereas Mff facilitates homeostatic fission (11). This is supported by numerous studies citing involvement of Fis1 in disease states, particularly neurodegenerative and cardiac disorders, as well as pulmonary hypertension and diabetic endothelial dysfunction (83–90). Further, treatment with a variety of chemical-based cellular stressors increases Fis1-Drp1 interactions by co-immunoprecipitation (35). It may be that Fis1 acts as a switch from homeostatic to mitophagic fission, signaling to Drp1 which portions of mitochondria need to be removed.

Ultimately, our results are consistent with a model in which human Fis1 has maintained a direct interaction with Drp1 via highly conserved residues to promote mitochondrial fission, resolving a long-standing question in the mitochondrial dynamics field. Further, this interaction is negatively regulated by the Fis1 N-terminal arm, suggesting the human and yeast proteins are more similar than previously thought (**Figure 6**). Similar to yeast where the Mdv1p adaptor is required(44, 91, 92), Fis1 and Drp1 alone are likely insufficient for mitochondrial fission and likely require other factors. Unlike the yeast system, perhaps Fis1’s interaction with Drp1 promotes mitophagic fission based on the work of others (11, 35), however, the studies presented here did not examine whether Fis1 or the Fis1 variants impact mitophagic and/or housekeeping fission. Our results suggest that Fis1 might sequester dimeric Drp1 poised for productive assembly. It may be that Fis1 is functioning in a similar manner to Mff which appears to preferentially recruit the dimeric species of Drp1 (93). Regardless, small changes to the Fis1 sequence result in a diverse spectrum of network phenotypes, suggesting that Fis1 is poised through its regulatory arm to sense environmental cues that can govern mitochondrial morphology.

**Figure 6.**
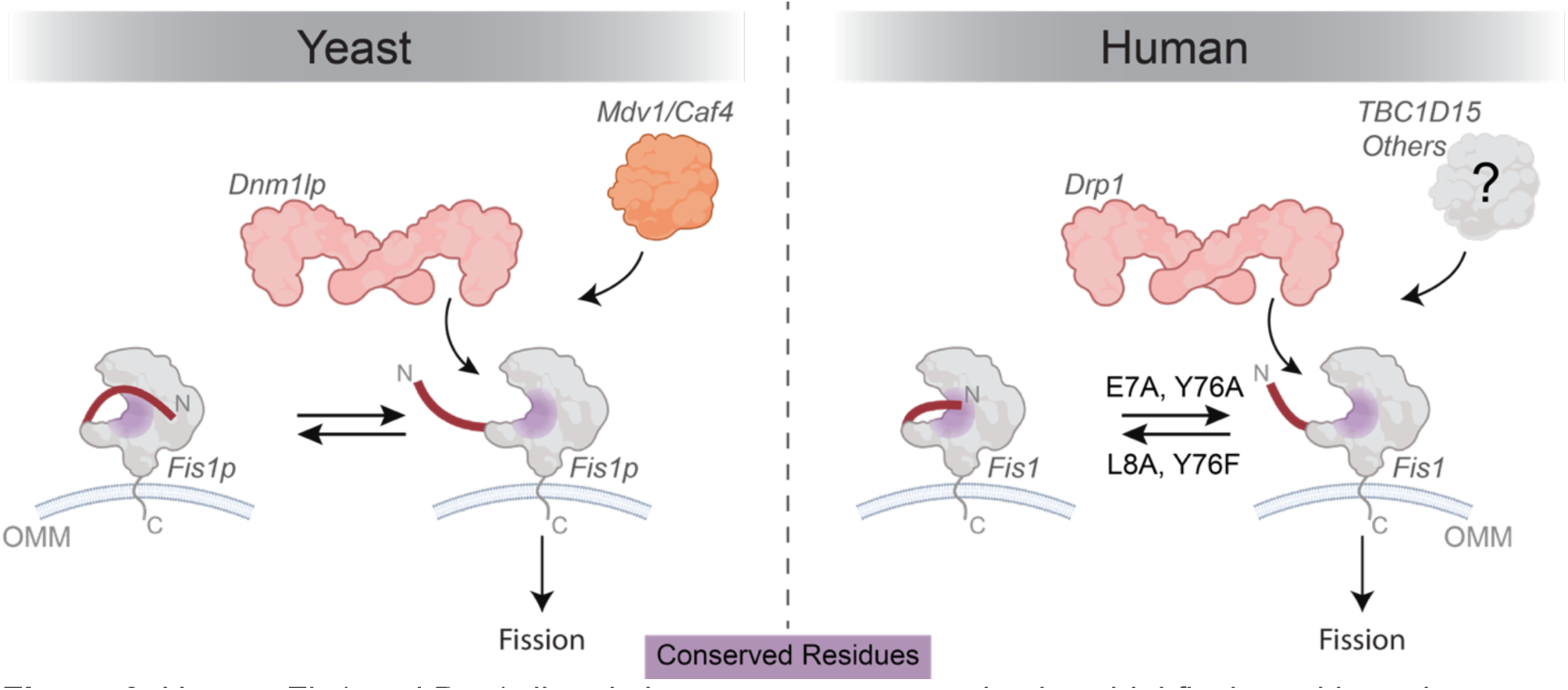
Human Fis1 and Drp1 directly interact to promote mitochondrial fission, akin to the yeast system. This interaction is governed by intramolecular interactions between the Fis1 arm and the protein’s conserved surface. Fis1 E7A and Y76A variants induce increased mitochondrial fragmentation, which may result from modulating Fis1 arm position into a more open, active conformation. In yeast, mitochondrial fission due to Fis1-Drp1 interactions (Fis1p-Dnm1p) relies on Mdv1/Caf4. Fis1-Drp1 mediated fission in humans likely relies on other proteins, such as TBC1D15.

## Experimental Procedures

### Protein Purification

#### Fis1 and Fis1ΔN

Recombinant Fis1^1-125^ or Fis1ΔN^9-125^ (WT and E7A) were expressed using pQE30 vectors as His_6_-Smt3-Fis1 fusion proteins in competent BL21 DE3 *Escherichia coli* also carrying the pREP4 plasmid. ^15^N-labeled protein samples were grown in 1L minimal media containing 3 g/L ^15^N ammonium chloride (Cambridge Isotope Laboratories, Tweksbury, MA) and unlabeled protein samples were grown in 1L super broth media. Bacteria were grown and protein was purified using nickel affinity chromatography as previously described (94) with the following changes: cells were lysed by sonication (four 30-second bursts with a 30-second rest period in between) and Buffer A consisted of 20 mM HEPES, pH 7.4, 500 mM NaCl, 40 mM Imidazole, 0.02% sodium azide. Samples were incubated with recombinant ULP1, the yeast SUMO protease, overnight at 4 °C (1:500 v/v) to remove the His_6_-Smt3 expression tag, leaving only native residues. The expression tag was then removed from the recombinant protein of interest by reverse nickel affinity chromatography. Samples were dialyzed into the Fis1 NMR buffer (20 mM HEPES, pH 7.4, 175 mM NaCl, 1 mM DTT, 0.02% sodium azide) using 6-8 kDa dialysis tubing (Repligen) at 4 °C overnight, undergoing at least one buffer exchange. Protein concentration was determined using the absorbance at 280 nm and the theoretical molar extinction coefficient determined from each constructs primary sequence. Purified protein samples were stored at 4 °C until use.

#### Drp1

Recombinant Drp1 isoform 1 was expressed and purified as previously described (56). Briefly, harvested cells were lysed using an EmlusiFlex C3 homogenizer (Avestin) at 15,000 p.s.i. and protein lysate was clarified via centrifugation at 15,000 rpm and 4C for 45 minutes using a JA-20 fixed-angle rotor in a Beckman J2-21 centrifuge. The clarified cell lysate was applied to a nickel affinity column (GE Healthcare; Sepharose high performance beads) equilibrated in Buffer A and purified by FPLC. Eluted protein was dialyzed into assay buffer (20 mM HEPES, pH 7.4, 150 mM KCl, 2.0 mM MgCl2, 1 mM DTT, 0.02% sodium azide) overnight at 4C using 6-8 kDa molecular weight cut-off dialysis tubing (Repligen) and the expression tag was simultaneously removed using recombinant TEV protease (∼1:20 ratio mg/mg). The remaining TEV protease was then removed using a reverse nickel chromatography step. Protein concentration was determined using the same approach as the Fis1 constructs. Purified protein was flash frozen in liquid nitrogen in single use aliquots consisting of 50-100 µL volume and stored at -80 °C until use. Protein was thawed on ice when needed. Purified Fis1 and Drp1 constructs were determined to be greater than 90% pure by SDS-PAGE analysis using a BioRad Mini-PROTEAN TGX Stain-Free gradient gel (4-20%).

#### Peptide Synthesis

The Fis1 N-terminal arm peptide (MEAVLNEL) with N-terminal acetylation and C-terminal amidation were purchased from GenScript who determined the peptide to be >95% pure by HPLC. The peptide was resuspended in 1.5 mL of 50 mM ammonium bicarbonate and required bath sonication to fully resuspend in solution. The peptide was then frozen at -80 °C in a 15 mL conical tube and lyophilized overnight to remove residual trifluoroacetic acid. The lyophilized peptide was then resuspended once more in ammonium bicarbonate and lyophilized as stated above. The peptide was weighed using an analytical balance and resuspended in Fis1 NMR buffer to the target concentration.

#### Thermal Shift Assays using NanoDSF

Protein unfolding as a function of temperature was measured by intrinsic protein fluorescence at 330 nm and 350 nm, and simultaneously by changes in light scattering, using a Prometheus NT.48 (NanoTemper, Germany) as previously described (33). Fis1^1-125^, Fis1ΔN^9-125^, and Fis1^1-125^ E7A were prepared at a final concentration of 30 µM in Fis1 NMR buffer (see above). A titration series to evaluate Fis1ΔN-Drp1 binding was performed so that the concentration of Drp1 was held constant at 30 µM as the amount of Fis1 or Fis1ΔN increased from 0-30 µM. Samples containing both Fis1ΔN and Drp1 were buffer matched so that each sample contained an equivalent amount of both Fis1 and Drp1 buffers to rule out buffer-dependent effects. Approximately 10 µL of each sample was loaded into Prometheus NT.48 Series nanoDSF high sensitivity capillaries (NanoTemper, Germany) and melting scans were performed using the Pr.ThermControl software (NanoTemper, Germany) with a temperature range of 25 °C to 95 °C and a temperature increase of 1 °C/minute. Melting temperatures (T_m_) were calculated from the temperature at the maximum value of the first derivative of the fluorescence signal at 330 nm. Data were imported into RStudio (4.2.2) (95) using readxl (96) and visualized using ggplot2 (3.3.3) (97) and RColorBrewer (98) as boxplots with temperature shown on the y-axis and protein constructs/concentration represented on the x-axis.

### Microscale Thermophoresis

#### Cy5-labeling of protein

Fis1ΔN and Drp1 were covalently modified at methionine residues in a two-step process using propargyl oxaziridine to generate a sulfimide conjugate to methionine containing a terminal alkyne and then using copper click chemistry to attach a Cy5-azide fluorophore(99, 100). To perform the labeling of Fis1ΔN(residues 9-125) that labels the only methionine, Met118, 10-fold molar excess propargyl oxaziridine was added to Fis1ΔN in Fis1 NMR buffer, vortexed to mix, and incubated at room temperature for 15 minutes. This was then applied to a PD10 column equilibrated in Fis1 NMR buffer and 1 mL fractions were collected. Fractions containing protein were concentrated to ∼0.25 mL using a 3 kDa concentrating tube and final protein concentration was determined using the absorbance at 280 nm and the protein’s theoretical extinction coefficient. HEPES and Cy5-azide dye (Click Chemistry Tools) were added to the protein sample at a final concentration of 25 mM and 350 µM, respectively. A final concentration of 1.25 mM BTTAA (2-(4-((Bis((1-(tert-butyl)-1H-1,2,3-triazol-4-yl)methyl)amino)methyl)-1H-1,2,3-triazol-1-yl)acetic acid, Click Chemistry Tools) and CuSO_4_ (250 µM final concentration) were combined and mixed until a light blue color was observed. Sodium ascorbate (at 12.5 mM final concentration) was then added to this mixture and vortexed until the solution turned clear. This mixture containing the BTTAA, CuSO_4_, and sodium ascorbate was then added to the protein sample and incubated in the dark at room temperature for 10 minutes. The sample was then applied to a PD10 column equilibrated in Fis1 NMR buffer and 1 mL fractions were collected as above. Fractions containing labeled protein were pooled and concentrated using a 3 kDa molecular weight cut-off concentrating tube to a final volume of ∼750 µL. Protein concentration and the degree of labelling were calculated using Equations 1 and 2 and a correction factor of 0.05 for Cy5 at 280nm. For Fis1ΔN (1 methionine) and Drp1 (15 methionines), the degree of labeling was less than 1 Cy5 per protein molecule. Cy5-protein samples were diluted to 10 µM in MST buffer (Cy5-Fis1ΔN: 20 mM HEPES, pH 7.4, 175 mM NaCl, 1 mM DTT, 0.05% TWEEN-20, 0.02% sodium azide; Cy5-Drp1: 20 mM HEPES, pH 7.4, 150 mM KCl, 2 mM MgCl2, 1 mM DTT, 0.05% TWEEN-20, 0.02% sodium azide), aliquoted into 20 µL samples, and frozen at -80 °C until needed.

Cy5-Fis1ΔN and Cy5-Drp1 were thawed on ice in the dark and then centrifuged using a tabletop centrifuge to remove aggregates. Cy5-Fis1ΔN and Cy5-Drp1 were then diluted to 80 nM or 200 nM working stocks in either Fis1 MST Buffer or Drp1 MST Buffer, respectively. For the Cy5-Drp1 titrations with Drp1 in the presence of Fis1ΔN, a stock of 80 nM Cy5-Drp1 and 2x the final concentration of Fis1ΔN were created and incubated in the dark for 1 hour at room temperature prior to titration set-up. A serial dilution of the unlabeled ligand of interest was performed in PCR tubes using ligand buffer (Drp1: 20 mM HEPES, pH 7.4, 150 mM KCl, 2 mM MgCl2, 1 mM DTT, 0.02% sodium azide; Fis1ΔN: 20 mM HEPES, pH 7.4, 175 mM NaCl, 1 mM DTT, 0.02% sodium azide) to create a total of 16 samples, each containing 10 µL of ligand in ligand buffer. 10 µL of Cy5-labeled protein (Cy5-Fis1ΔN = 40 nM final; Cy5-Drp1 = 80 nM final) was then mixed with each ligand only sample in a 1:1 ratio for a total volume of 20 µL. Samples were then briefly spun-down using a tabletop centrifuge and incubated in the dark at room temperature for one hour prior to being loaded into Monolith Standard Capillaries (NanoTemper, Germany) and microscale thermophoresis measurements were performed using the MO.Control Software with a medium laser power and fluorescence auto-detection. Changes in the relative normalized fluorescence (Cy5-Fis1ΔN and Cy5-Drp1 + Fis1ΔN) or changes in initial fluorescence intensity (Cy5-Drp1) as a function of ligand concentration were plotted in R and fit to a four-parameter log-logistic model to determine apparent binding affinities ± error of the fit. Data representative of n=3. Cy5-Drp1 titration data in the presence of Fis1ΔN was fit to an isodesmic assembly model in Igor Pro (WaveMetrics) to determine Drp1 self-affinity. These data were then converted to fraction bound assuming 100% bound state at the highest concentration of Fis1ΔN and fit to a 1:1 binding model to determine an apparent affinity of Drp1 for Fis1ΔN.

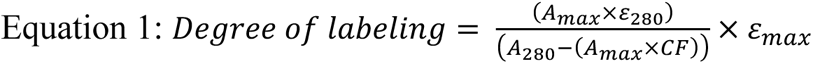

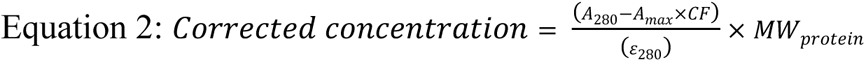

Where A_max_ = A_649_, CF = 0.05, ε_max_ = 250,000 M^-1^cm^-1^ for Cy5 as provided in the product specifications by Click Chemistry Tools. MW_Fis1ΔN_ = 13,584 daltons and MW_Drp1_ = 81,877 daltons.

### NMR Spectroscopy

#### 1H, 15N HSQC

^1^H, ^15^N HSQC spectra were collected on 50 µM ^15^N-Fis1^1-125^, ^15^N-Fis1ΔN^9-125^ in 10% D_2_O (Fis1 NMR Buffer: 20 mM HEPES, pH 7.4, 175 mM NaCl, 1 mM DTT, 0.02% sodium azide) ± Drp1 (isoform 1 or 5) or the N-terminal arm peptide at the indicated concentrations at 25 °C using a Bruker Avance II 600 MHz spectrometer. Samples without ligand were buffer matched using an equivalent volume of ligand buffer (Drp1 dialysate buffer: 20 mM HEPES, pH 7.4, 150 mM KCl, 2 mM MgCl2, 1 mM DTT, 0.02% sodium azide or peptide: Fis1 NMR buffer) to account for potential spectral changes due to buffer conditions. ^1^H,^15^N HSQC spectra collected on ^15^N-Fis1^1-125^ E7A under similar conditions. The spectrometer was equipped with a z-axis gradient cryoprobe and SampleJet autosampler, which allowed for automatic tuning, shimming, and data collection of samples. Experiments were comprised of 8 scans with 1024 and 300 complex points in the ^1^H and ^15^N dimensions, respectively. The ^15^N-Fis1ΔN + Drp1 titration spectra were processed using NMRPipe as previously described (33) and chemical shift intensity changes were measured using TitrView and CARA software. Chemical shift values for all other ^1^H, ^15^N experiments were measured using CCP4 NMRAnalysis (CcpNmr version 2.5.2) on NMRBox (101) via the snap assignment feature. Chemical shift values for each residue were then used to determine chemical shift perturbations as previously described (102) using Equation 3 which were then plotted in RStudio (4.2.2) as a function of protein residue number using ggplot2. Δδ and ΔIntensity values were displayed on the NMR structure of Fis1 (PDB: 1PC2) truncated to reflect the constructs used in their respective experiments using PyMOL (Schrödinger).

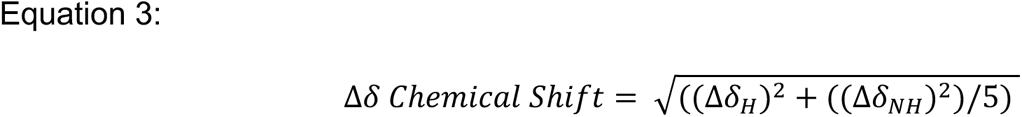

An apparent binding affinity for the N-terminal arm peptide was determined using TREND analysis(103), which performs a principal component analysis of each spectrum within its respective titration series, with an output of normalized principal component 1 values (PC1). The PC1 values were plotted in RStudio using ggplot2 as a function of ligand concentration and fit to a one-site binding model (Equation 4) in which protein concentration was held constant. Apparent binding affinities for Drp1 were determined by plotting the ΔIntensity values of each Fis1ΔN residue as a function of Drp1 concentration and fitting the data to Equation 4. Spectral overlays were generated using XEASY and Adobe Illustrator.

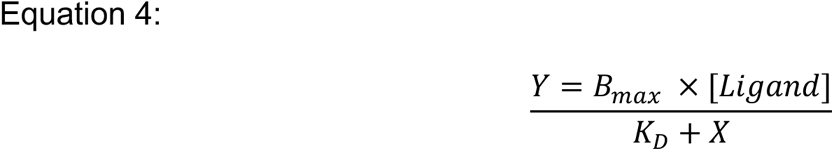

#### Drp1 GTP Hydrolysis Assays

Drp1 (isoform 3) hydrolytic activity was measured using a Malachite Green-based colorimetric assay to detect free phosphate in solution. Drp1 isoform 3 (500 nM final concentration) was prepared in 10x Assay Buffer (final concentrations: 20 mM HEPES, pH 7.4, 50 mM KCl, 2 mM MgCl_2_, 1 mM DTT) ± Fis1 or Fis1ΔN (final concentration 25 µM). Drp1 only samples were prepared with an equivalent amount of Fis1 buffer (20 mM HEPES, pH 7.4, 175 mM NaCl, 1 mM DTT, 0.02% sodium azide) to account for potential effects due to differing buffer compositions. Phosphate standards (0, 4, 8, 12, 16, 24, 32, and 40 µM) were created in ddH2O from a 1 mM potassium phosphate stock. Samples were incubated 1 hour at room temperature to ensure binding interacts were at equilibrium prior to aliquoting 72 µL of Drp1 containing reaction mix into wells of a flat-bottom 96-well plate (Corning Costar). A total of 5 wells were used per replicate to account for 5 time points (0, 10, 20, 30, and 40 minutes). The initial timepoint (0 minutes) was quenched using 80 µL 2x EDTA, pH 8.0 (final concentration 25 mM) prior to starting the assay. To start the reactions, 8 µL of 10x GTP (1 mM final concentration) was added to each sample mixture using a multi-channel pipette. Plate was placed in a 37 °C plate shaker (600 RPM, Multi-Therm Shaker) and reaction was allowed to proceed for 40 minutes. Every ten minutes, an additional well was quenched using 80 µL of 2x EDTA, pH 8.0. After reactions were completed, 40 µL of Malachite Green Detection Solution (5 mL Malachite Green dye solution, 1.25 mL of 15.24% weight/volume ammonium molybdate tetrahydrate, and 0.1 mL 11% weight/volume TWEEN-20) was added to the samples and phosphate standards and allowed to incubate at RT for 2 minutes. Malachite Green dye solution prepared from 0.44 g Malachite Green hydrochloride, 300 mL ddH_2_O, and 60 mL concentrated H_2_SO_4_. After this, 20 µL of 34% sodium citrate was added to each well (final concentration of 3.1%) to adjust pH to prevent nonenzymatic hydrolysis of remaining GTP. Samples incubated an additional 10 minutes at RT prior to reading Abs620 nm values using a FlexStation plate reader (Molecular Devices). Data were imported into R.Studio where the phosphate standards were used to generate a phosphate standard curve. The standard curve was then used to convert experimentally derived Abs620 nm values into phosphate concentrations. Phosphate release over time was then used to calculate GTPase rates for each sample. Data were visualized using the ggplot2 package.

#### Cell culture and Transfection

Human RPE cells, wild-type or Fis1*^-/-^*generated via CRISPR/Cas9, were cultured without antibiotics and transfected as previously described (33). Briefly, cells were grown in Dulbecco’s modified Eagle medium (DMEM)-F12 (Thermo Fischer Scientific) with 10% fetal bovine serum (Gemini). Cells were plated in No. 1.5 glass bottom 24-well dishes (Cellvis) and transfected approximately 24 hours after plating. Plasmid DNA was added to Opti-MEM and mixed by vortexing before adding Avalanche-Omni to the DNA:Opti-MEM mixture and vortexing for 5 seconds. The transfection complexes were then incubated at room temperature for 15 minutes and added dropwise into each well. Cells were then incubated overnight and subsequently processed for immunofluorescence.

#### Immunofluorescence Staining and Imaging

Cells were prepared for immunofluorescence staining using the methods described in (33). To reduce nonspecific binding, a blocking solution of 3% BSA (w/v)/0.3% Triton X-100 in PBS was used following permeabilization, as well as in each antibody incubation step. To minimize antibody cross reactivity in experiments with dual-labeling, immunofluorescence steps were performed sequentially with Drp1 staining first, followed by Fis1. Reagents and concentrations described in more detail in the supporting information. Cells were stored and imaged in PBS using a 60x oil objective. Representative confocal images were acquired using a spinning disc confocal microscope (Nikon Eclipse Ti2-E microscope) at 0.3 µm z-slices and 0.11 µm/pixel resolution and processed using ImageJ.

#### Protein Colocalization, Fis1 intensity, and MitoGraph Analysis

Images were prepared for colocalization, fluorescence intensity, and mitochondrial network morphology analysis using the methods in (33). Per the protocol, ImageJ macros were used to create single channel/single cell z-stack images based on regions of interest (ROIs) generated in ImageJ. Protein colocalization between endogenous Drp1 and mitoYFP was determined using a Pearson’s correlation analysis via the coloc 2 function on all z-slices. Fis1 intensity was measured using the ROIs from maximum intensity projection image stacks.

For a detailed description of the MitoGraph analysis methods, please see (33). Briefly, PNG files of individually cropped cells underwent mitochondrial segmentation based on the mitoYFP signal and were then visually screened for accurate segmentation. A denoising step was performed. R scripts were used to extract a variety of metrics, including total connected components, PHI, average edge length, total edges, total nodes, 3-way and 4-way junctions, and the MitoGraph Connectivity Score, from the GNET files produced by MitoGraph. Data were imported into RStudio and the Pearson’s scores, MitoGraph metrics, and Fis1 intensity analysis were merged into a single dataset. Data were visualized using ggplot2 as boxplots. Statistical analysis was performed using an ANOVA with post-hoc Tukey’s. MitoGraph can be downloaded free of charge at https://github.com/vianamp/MitoGraph. R-scripts used for MitoGraph analysis are available at https://github.com/Hill-Lab/MitoGraph-Contrib-RScripts.

## Data Availability

All R scripts used for data analysis and visualization are available for download at https://github.com/Hill-Lab/. Raw data is available upon request from Blake Hill at.

## Supporting Information

This article contains supporting information.

## Supporting information

Supporting Informatino

## Acknowledgments

We would like to thank Drs. F.C. Peterson and Dr. R. Keyes of the Medical College of Wisconsin Program in Chemical Biology for the click-chemistry reagents and guidance. In addition, we would like to thank the members of the Hill laboratory for their discussions and support.

## Funding and additional information

This project was supported by the following National Institutes of Health grants: R01GM067180 (to RB Hill) and TL1TR001437 and T32GM080202 (to KA Nolden). The content is solely the responsibility of the authors and does not necessarily represent the official views of the National Institutes of Health.

## Conflicts of interest

RBH and KAN have a financial interest in Cytegen, a company developing therapies to improve mitochondrial function. However, neither the research described herein was supported by Cytegen, nor was in collaboration with the company.

