## Supporting Informatino for "Human Fis1 directly interacts with Drp1 in an evolutionarily conserved manner to promote mitochondrial fission"

**This PDF file includes:**

**Supporting Information Figure Legends**

**Figure S1.** Highly conserved Fis1 residues localize to the protein's concave surface.
**Figure S2.** Fis1, ΔN, and E7A directly interact with Drp1 with the same affinity.
**Figure S3.** Fis1 specifically binds to Drp1 in an isoform independent manner.
**Figure S4.** Fis1 does not affect Drp1 GTP hydrolysis in solution.
**Figure S5.** Representative confocal images of arm alanine scan experiment.
**Figure S6.** MitoGraph Score metrics for alanine scanning of Fis1 arm**.**
**Figure S7.** Fis1 mean intensity vs. MitoGraph or Drp1-mitochondrial colocalization for the Fis1 arm alanine scan experiment.
**Figure S8.** Correlation between Drp1-mitochondrial colocalization and PHI.
**Figure S9.** Fis1 E7A binds Drp1 by 2D NMR.
**Figure S10.** Representative confocal images of Fis1 concave surface mutation experiments.
**Figure S11.** MitoGraph Score metrics for Fis1 concave surface mutations experiments.
**Figure S12.** Fis1 mean intensity vs. MitoGraph or Drp1-mitochondrial colocalization for the Fis1 concave surface mutations experiment.
**Figure S13.** Fis1ΔN binds an assembly deficient Drp1 variant with similar affinity as wild-type.

**Table S1.** Table of reagents
**Table S2.** Consurf score for Fis1$^{1\text{-}125}$

### Supporting Information Figure Legends

**Figure S1. Highly conserved Fis1 residues localize to the protein's concave surface.** The cytoplasmic domain of Fis1 adopts a TPR fold comprised of two TPR repeats flanked by helices. The TPR fold creates a concave surface (left) and a convex surface (right). To determine evolutionarily conserved residues of Fis1, a ConSurf analysis was performed on the human Fis1 NMR structure (PDB:1PC2) (Table S2) and the resulting scores displayed on the surface representation of Fis1 (PDB: 1PC2$^{1\text{-}125}$) in a gradient from white to blue where a darker blue color correlates to a higher degree of conservation.

**Figure S2. Fis1, ΔN, and E7A directly interact with Drp1 with the same affinity.** ***A***. $\Delta F_{norm}$ values of 40 nM Cy5-Fis1ΔN in the presence of increasing unlabeled Drp1 as determined by microscale thermophoresis and fit to a single-site binding isotherm gives an apparent $K_D$ = 15 ± 5 μM. $\Delta F_{norm}$ values normalized to a 0-1 scale to allow for comparisons and averaging between multiple experiments (n=3). One experiment was not averaged due to differing ligand concentrations (shown without error bars). Error bars = SD of the two experiments performed with equivalent Drp1 concentrations. *B.* ΔFluorescence of 80 nM Cy5-Drp1 titrated with increasing Fis1. Data fit as in (A) indicating an apparent $K_D$ = 13 ± 3 μM. N=1. *C.* dFluorescence (330 nm) of 30 μM Drp1 titrated with increasing Fis1ΔN (0-30 μM) as a function of increasing temperature as determined by differential scanning fluorimetry. *D.* Box plots depicting the thermal unmelting temperature ($T_m$) of 30 μM Drp1 ± 30 μM Fis1ΔN, Fis1, and Fis1 E7A. $T_m$ values determined as the temperature corresponding to the first derivative of the maximum fluorescence value (n=2 for Fis1ΔN ± Drp1, n=3 for all others). *E.* Full $^{1}H$, $^{15}N$ spectral overlays of 50 μM $^{15}N$-Fis1ΔN in the presence of increasing Drp1 (0-85 μM). *F.* $^{1}H$, $^{15}N$ spectral overlay of 50 μM $^{15}N$-Fis1 ± 140 μM Drp1 with crosspeak intensity loss indicative of binding.

**Figure S3. Fis1 specifically binds to Drp1 in an isoform independent manner.** *A.* $^{1}H$, $^{15}N$ spectral overlays of 50 μM $^{15}N$-Fis1ΔN ± 400 μM BSA without chemical shift perturbation or intensity changes, suggesting no binding. *B.* $^{1}H$, $^{15}N$ spectral overlays of 50 μM $^{15}N$-Fis1ΔN ± 83 μM Drp1 isoform 1 (left) and 83 μM Drp1 isoform 5 (right).

**Figure S4. Fis1 does not affect Drp1 GTP hydrolysis in solution.** *A.* GTPase activity of Drp1 in solution in the presence and absence of Fis1 or Fis1ΔN shown as phosphate released over duration of the assay. *B.* Data in A converted to rate of phosphate release which indicate no significant changes in Drp1 hydrolytic rates upon Fis1 or Fis1ΔN addition. N=2. Error bars = SD.

**Figure S5. Representative confocal images of arm alanine scan experiment.** Cells transfected with mito-YFP and either pcDNA, pcDNA-Fis1, or pcDNA-Fis1 arm point substitutions with boxes indicating the individual cells shown in Figure 3. Merged images show mito-YFP (green) and anti-Drp1 (magenta).

**Figure S6. MitoGraph Score metrics for alanine scanning of Fis1 arm .** Metrics calculated from single-cell z-stack images of mito-YFP transfected cells which were segmented using MitoGraph on the mito-YFP channel for the N-terminal arm alanine scanning experiment in Figure 3. PHI = fraction of total mitochondria occupied by a single, large mitochondrion called a connected component, average edge length = distance between branch points or length of individual mitochondrion, nodes = number of branch points or termini, edges = number of branches or individual mitochondrion, connected components = number of connected mitochondria in a cell, and average degree = based on nearest neighbor analysis identifying free ends and branch points).

**Figure S7. Fis1 mean intensity vs. MitoGraph or Drp1-mitochondrial colocalization for the Fis1 arm alanine scan experiment.** To determine relative expression levels of Fis1 and variants, Fis1 mean intensity from immunofluorescence was calculated from single cell z-stack images and compared to MitoGraph Connectivity Score (A) and Pearson's colocalization between mito-YFP and Drp1 (B) for each transfection condition for the N-terminal arm alanine scanning experiment in Figure 3. Data fit were fit to a least-squares linear regression with $R^2$ value of the fit shown in the top right of each plot.

**Figure S8. Correlation between Drp1-mitochondrial colocalization and PHI.** (*A, B*) Analysis of Drp1-mitochondrial localization in terms of Pearson score from Figures 3 (A, alanine scanning experiment) and 5 (B, conserved residue mutations) as a function of PHI score. Drp1-mitochondrial colocalization increases in cells with more highly clumped mitochondrial networks as indicated by a higher PHI score and Pearson score. Conversely, cells with lower PHI scores (indicative of more fragmented mitochondrial networks without clumping) correlates with lower Drp1-mitochondrial colocalization as indicated by a lower Pearson value. Gradient color indicates Fis1 expression in terms of Fis1 mean intensity.

**Figure S9. Fis1 E7A binds Drp1 by 2D NMR.** $^1$H, $^{15}$N HSQC spectral overlay of 50 µM $^{15}$N-Fis1 E7A ± 140 µM Drp1 with crosspeak intensity loss upon addition of unlabeled Drp1 indicative of binding.

**Figure S10. Representative confocal images of Fis1 concave surface mutation experiments.** Cells transfected with mito-YFP and either pcDNA, pcDNA-Fis1, or pcDNA-Fis1 point substitutions with boxes indicating the individual cells shown in Figure 5. Merged images show mito-YFP (green) and anti-Drp1 (magenta).

**Figure S11. MitoGraph Score metrics for Fis1 concave surface mutations experiments.** Metrics calculated from single cell z-stack images of mito-YFP transfected cells which were segmented using MitoGraph on the mito-YFP channel for the Fis1 point substitution experiment in Figure 5. See Figure S6 for definitions.

**Figure S12. Fis1 mean intensity vs. MitoGraph or Drp1-mitochondrial colocalization for the Fis1 concave surface mutations experiment.** Fis1 mean intensity calculated from single cell z-stack images compared to MitoGraph Connectivity Score (*A*) and Pearson's colocalization between mito-YFP and Drp1 (*B*) for each transfection condition for the Fis1 point substitutions experiment in Figure 5. Data fit to a linear equation with $R^2$ value of the fit shown in the top right of each plot.

**Figure S13. Fis1ΔN binds an assembly deficient Drp1 variant with similar affinity as wild-type.** $\Delta F_{norm}$ values of 40 nM Cy5-Fis1ΔN in the presence of increasing unlabeled Drp1 G401S, an assembly-deficient pathological Drp1 variant, fit to a single-site binding isotherm indicates Fis1ΔN has a similar affinity to an obligate-dimeric Drp1 in solution. N=1.

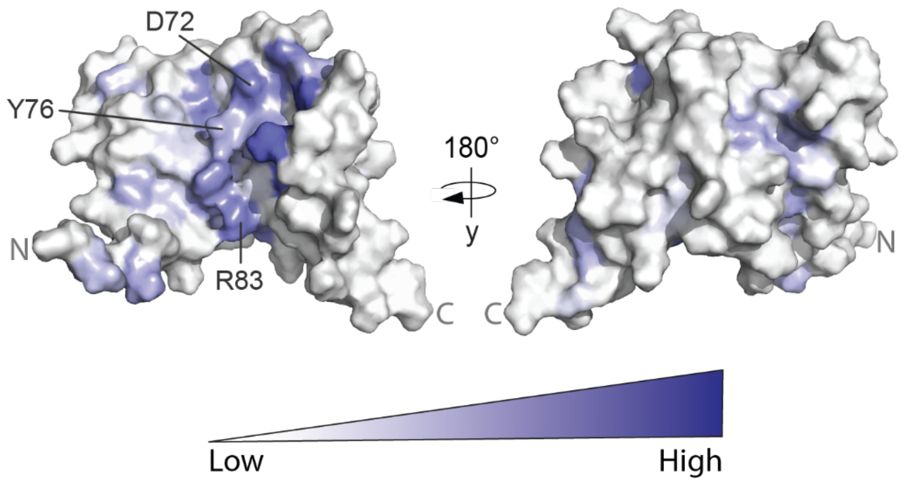


**Figure S1.**

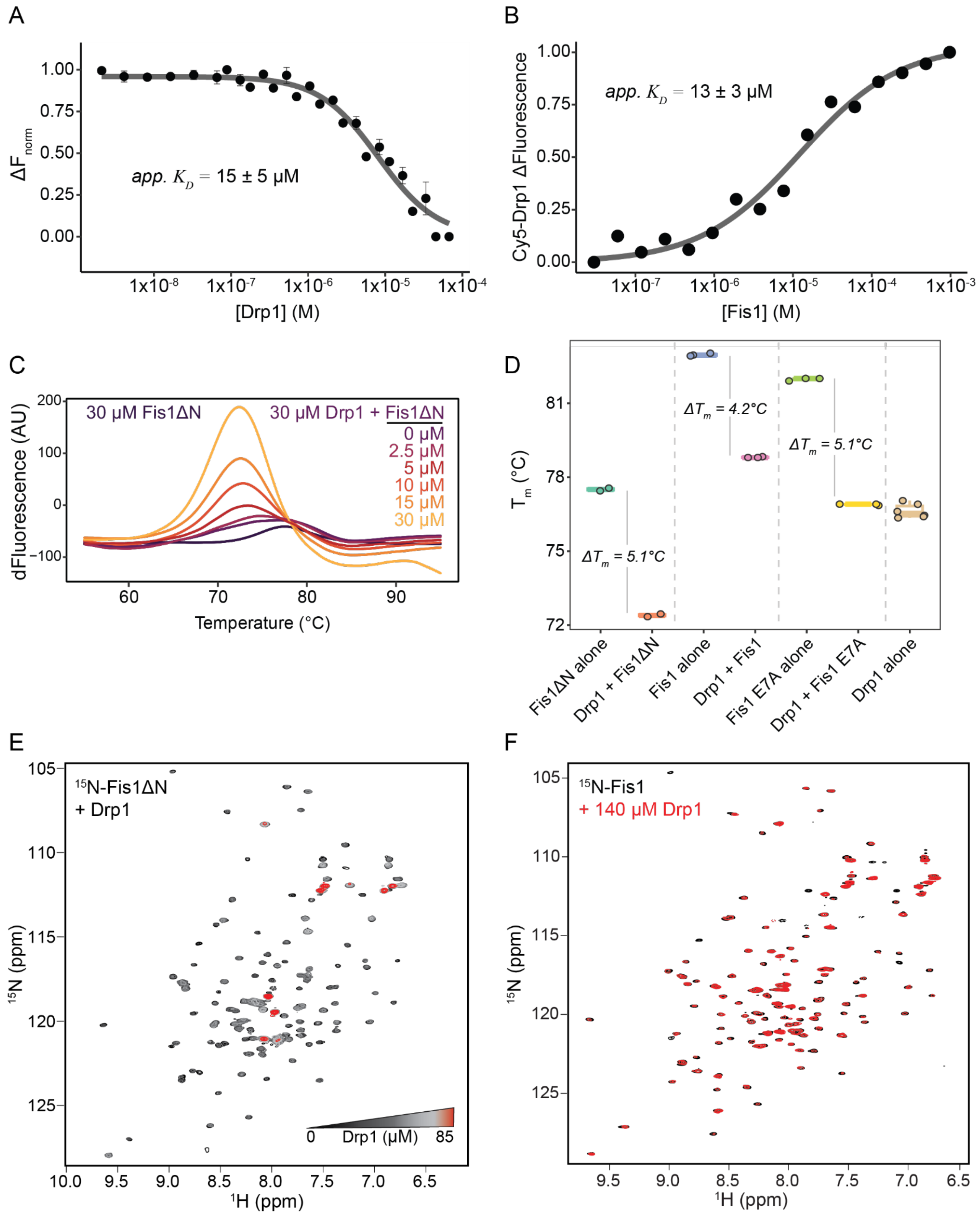
A
1.00
0.75
0.50
0.25
0.00
ΔF norm
app. K_D = 15 ± 5 μM
1x10-8
1x10-7
1x10-6
1x10-5
1x10-4
[Drp1] (M)
B
Cy5-Drp1 ΔFluorescence
app. K_D = 13 ± 3 μM
1x10-7
1x10-6
1x10-5
1x10-4
1x10-3
[Fis1] (M)
C
30 μM Fis1ΔN
30 μM Drp1 + Fis1ΔN
0 μM
2.5 μM
5 μM
10 μM
15 μM
30 μM
dFluorescence (AU)
200
100
0
−100
60
70
80
90
Temperature (°C)
D
81
78
75
72
T_m (°C)
ΔT_m = 4.2°C
ΔT_m = 5.1°C
ΔT_m = 5.1°C
Fis1ΔN alone
Drp1 + Fis1ΔN
Fis1 alone
Drp1 + Fis1
Fis1 E7A alone
Drp1 + Fis1 E7A
Drp1 alone
E
15N-Fis1ΔN
+ Drp1
0
Drp1 (μM)
85
15N (ppm)
1H (ppm)
F
15N-Fis1
+ 140 μM Drp1
15N (ppm)
1H (ppm)

**Figure S2.**

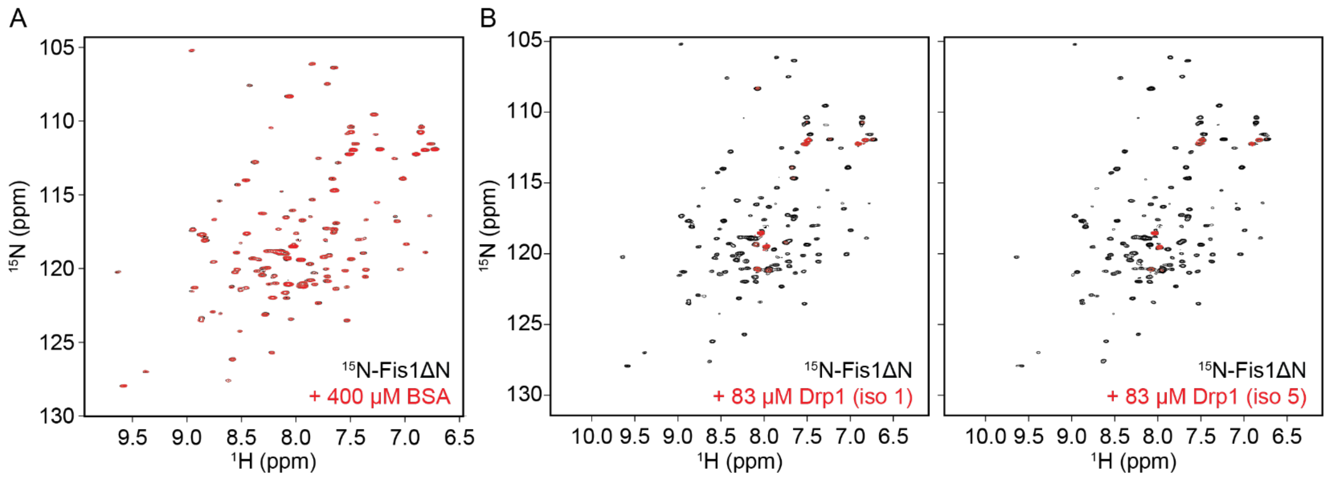
A
B
105
110
115
120
125
130
15N (ppm)
1H (ppm)
9.5 9.0 8.5 8.0 7.5 7.0 6.5
10.0 9.5 9.0 8.5 8.0 7.5 7.0 6.5
15N-Fis1ΔN
+ 400 μM BSA
+ 83 μM Drp1 (iso 1)
+ 83 μM Drp1 (iso 5)

**Figure S3.**

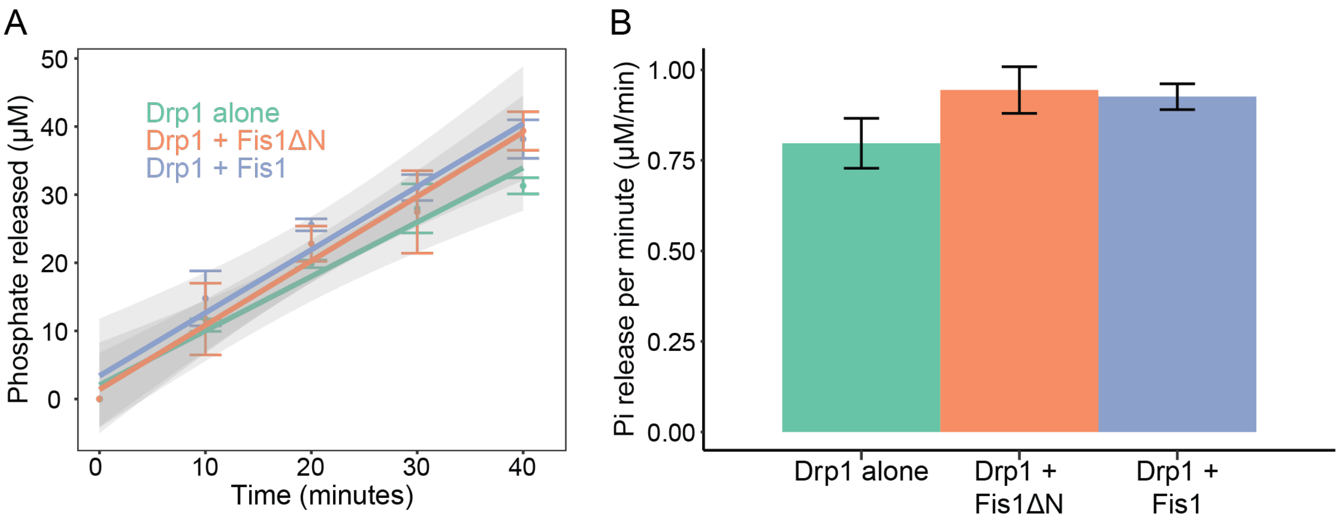
A
50
40
30
20
10
0
Phosphate released (μM)
Drp1 alone
Drp1 + Fis1ΔN
Drp1 + Fis1
0
10
20
30
40
Time (minutes)
B
Pi release per minute (μM/min)
1.00
0.75
0.50
0.25
0.00
Drp1 alone
Drp1 + Fis1ΔN
Drp1 + Fis1

**Figure S4.**

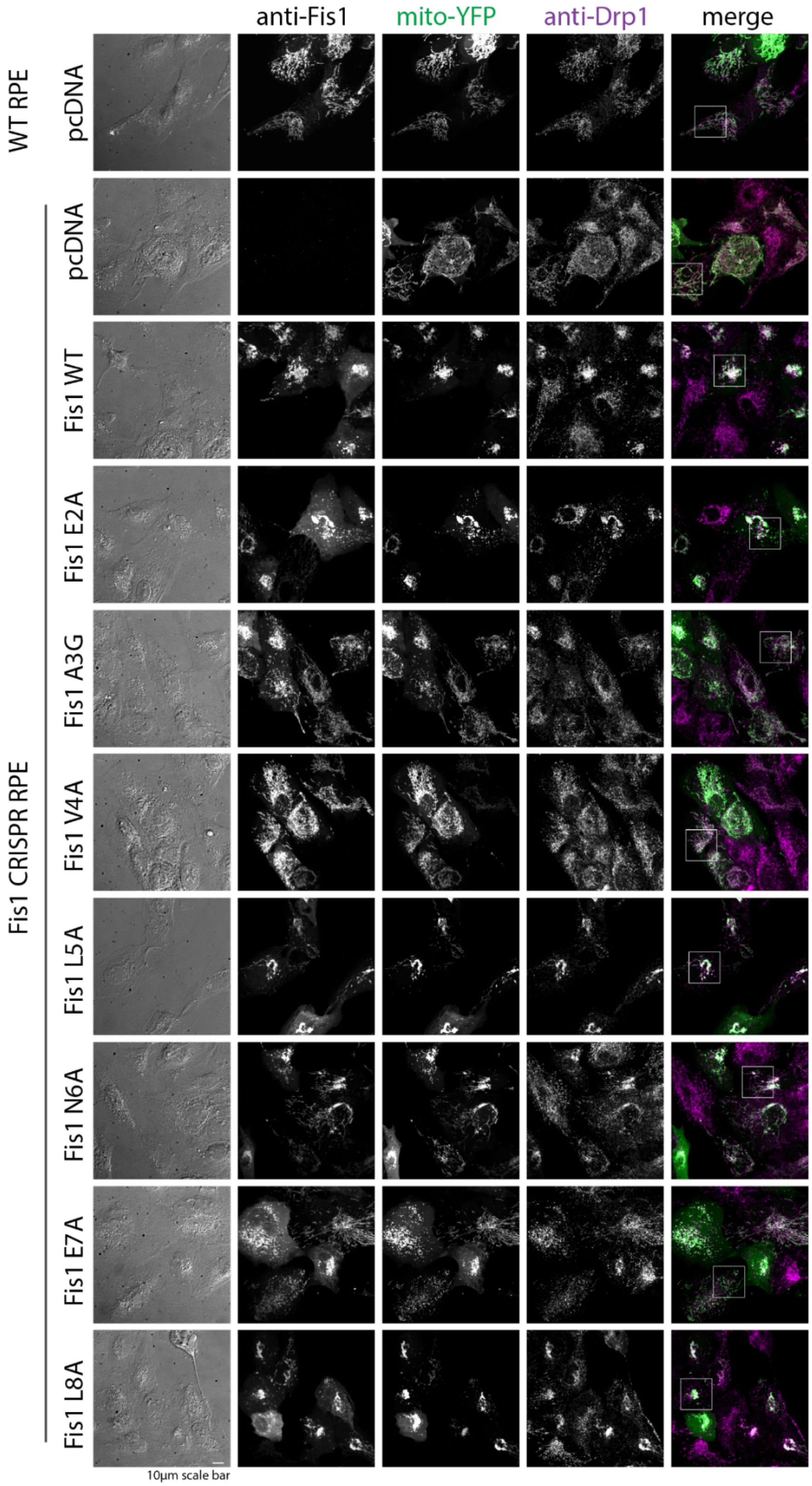
anti-Fis1
mito-YFP
anti-Drp1
merge
WT RPE
pcDNA
Fis1 CRISPR RPE
pcDNA
Fis1 WT
Fis1 E2A
Fis1 A3G
Fis1 V4A
Fis1 L5A
Fis1 N6A
Fis1 E7A
Fis1 L8A
10μm scale bar

**Figure S5.**

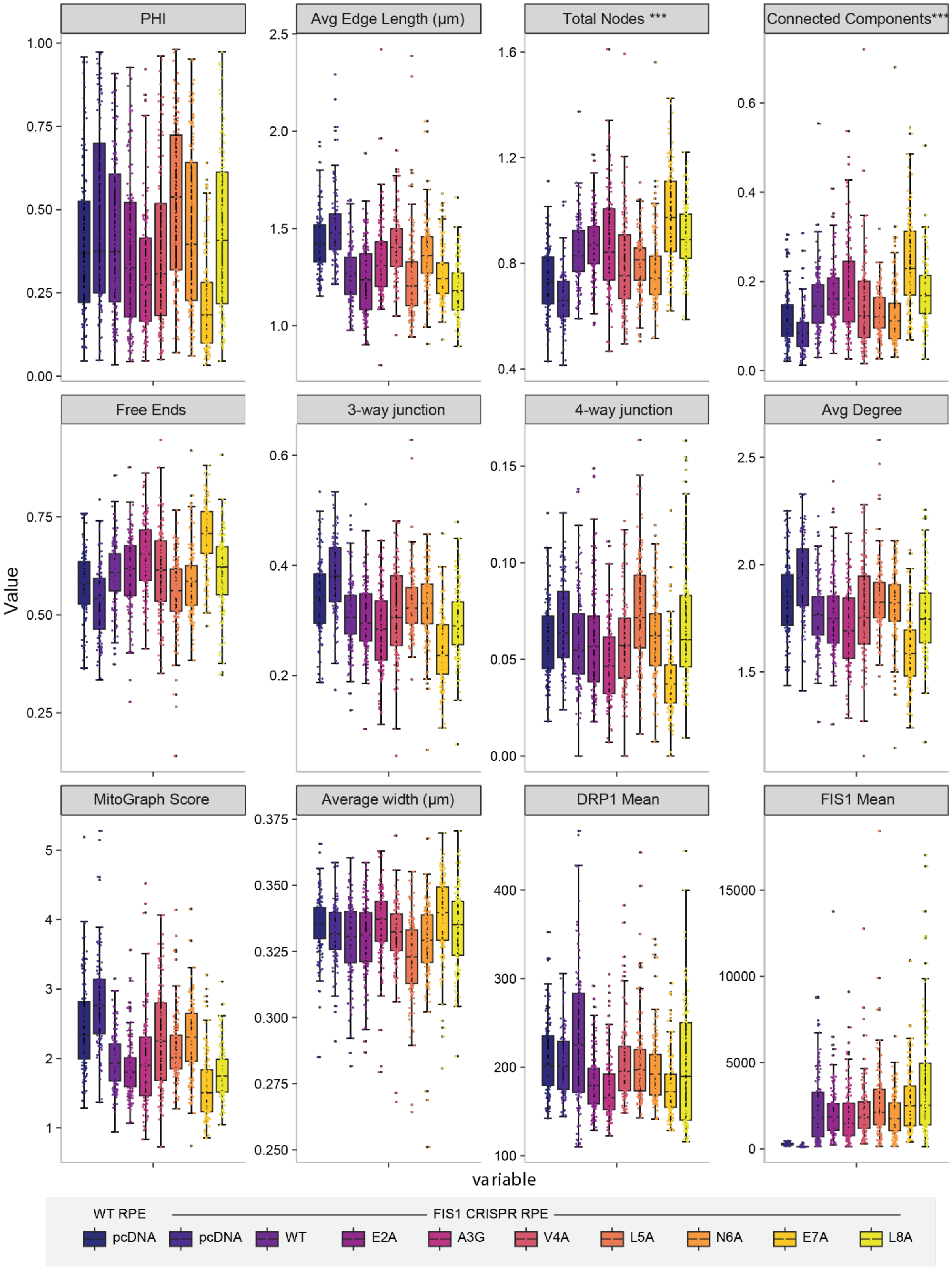

PHI
Avg Edge Length (μm)
Total Nodes ***
Connected Components***
Free Ends
3-way junction
4-way junction
Avg Degree
MitoGraph Score
Average width (μm)
DRP1 Mean
FIS1 Mean
Value
variable
WT RPE
FIS1 CRISPR RPE
pcDNA
pcDNA
WT
E2A
A3G
V4A
L5A
N6A
E7A
L8A


**Figure S6**.

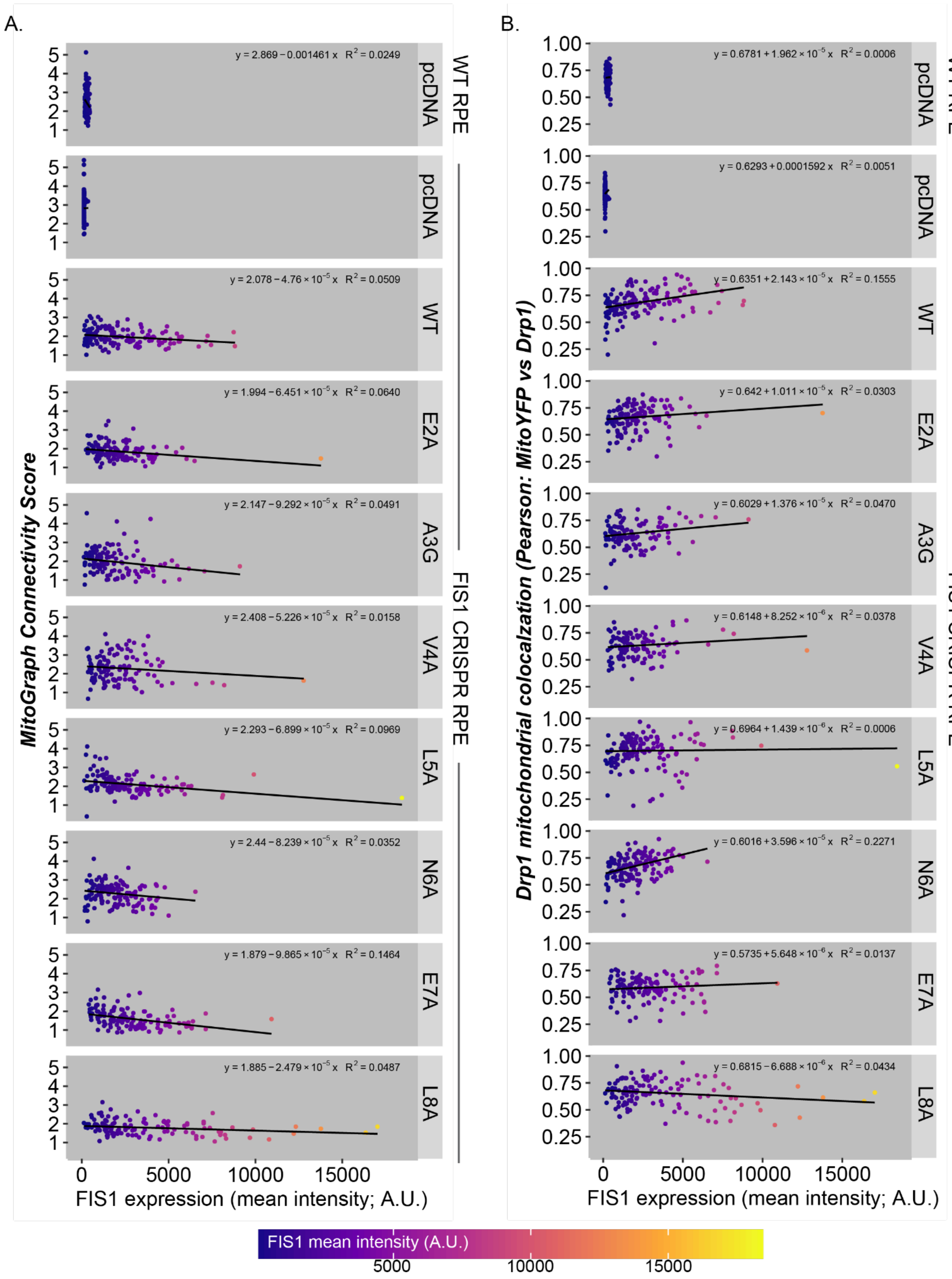
A.
B.
MitoGraph Connectivity Score
Drp1 mitochondrial colocalzation (Pearson: MitoYFP vs Drp1)
FIS1 expression (mean intensity; A.U.)
WT RPE
FIS1 CRISPR RPE
pcDNA
WT
E2A
A3G
V4A
L5A
N6A
E7A
L8A
y = 2.869 − 0.001461 x R2 = 0.0249
y = 2.078 − 4.76 × 10−5 x R2 = 0.0509
y = 1.994 − 6.451 × 10−5 x R2 = 0.0640
y = 2.147 − 9.292 × 10−5 x R2 = 0.0491
y = 2.408 − 5.226 × 10−5 x R2 = 0.0158
y = 2.293 − 6.899 × 10−5 x R2 = 0.0969
y = 2.44 − 8.239 × 10−5 x R2 = 0.0352
y = 1.879 − 9.865 × 10−5 x R2 = 0.1464
y = 1.885 − 2.479 × 10−5 x R2 = 0.0487
y = 0.6781 + 1.962 × 10−5 x R2 = 0.0006
y = 0.6293 + 0.0001592 x R2 = 0.0051
y = 0.6351 + 2.143 × 10−5 x R2 = 0.1555
y = 0.642 + 1.011 × 10−5 x R2 = 0.0303
y = 0.6029 + 1.376 × 10−5 x R2 = 0.0470
y = 0.6148 + 8.252 × 10−6 x R2 = 0.0378
y = 0.6964 + 1.439 × 10−6 x R2 = 0.0006
y = 0.6016 + 3.596 × 10−5 x R2 = 0.2271
y = 0.5735 + 5.648 × 10−6 x R2 = 0.0137
y = 0.6815 − 6.688 × 10−6 x R2 = 0.0434
FIS1 mean intensity (A.U.)
5000
10000
15000

**Figure S7.**

A.

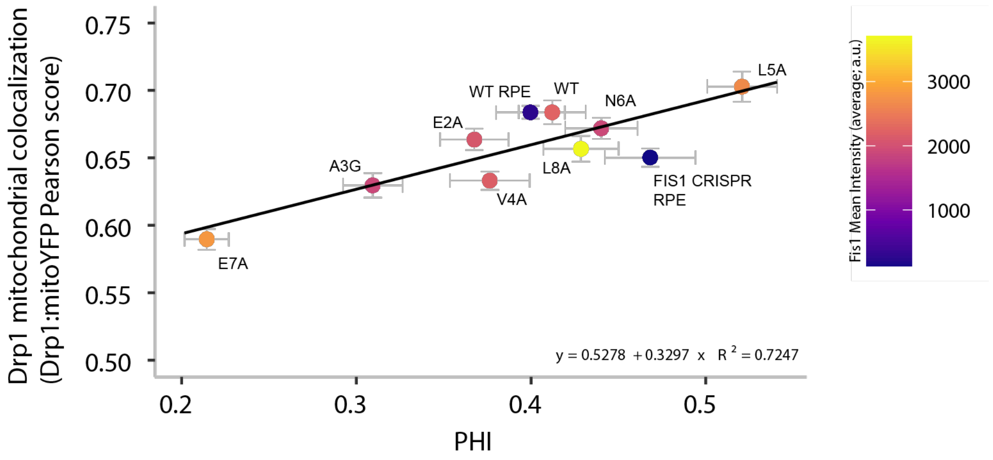

Drp1 mitochondrial colocalization
(Drp1:mitoYFP Pearson score)
0.75
0.70
0.65
0.60
0.55
0.50
0.2
0.3
0.4
0.5
PHI
WT RPE
WT
L5A
N6A
E2A
A3G
L8A
V4A
FIS1 CRISPR
RPE
E7A
y = 0.5278 + 0.3297 x R 2 = 0.7247
Fis1 Mean Intensity (average; a.u.)
3000
2000
1000


B.

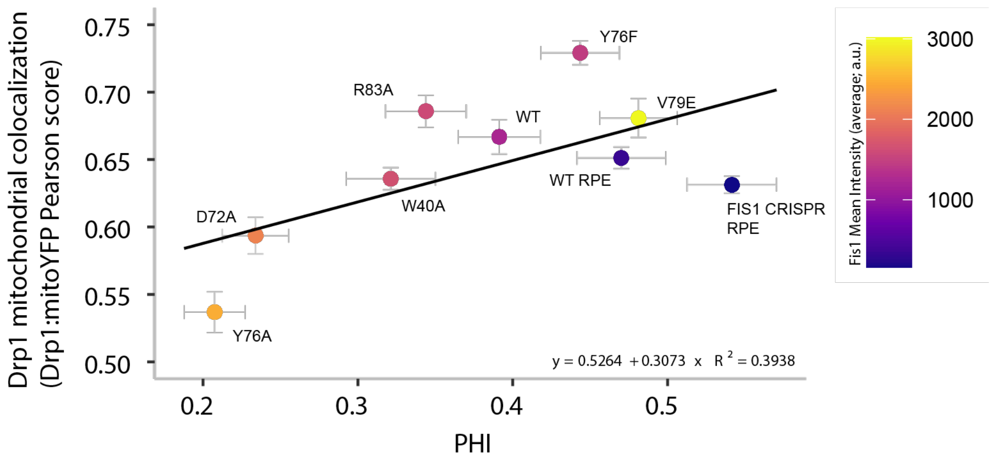

Drp1 mitochondrial colocalization
(Drp1:mitoYFP Pearson score)
0.75
0.70
0.65
0.60
0.55
0.50
0.2
0.3
0.4
0.5
PHI
Y76F
R83A
V79E
WT
WT RPE
W40A
D72A
FIS1 CRISPR
RPE
Y76A
y = 0.5264 + 0.3073 x R 2 = 0.3938
Fis1 Mean Intensity (average; a.u.)
3000
2000
1000


**Figure S8.**

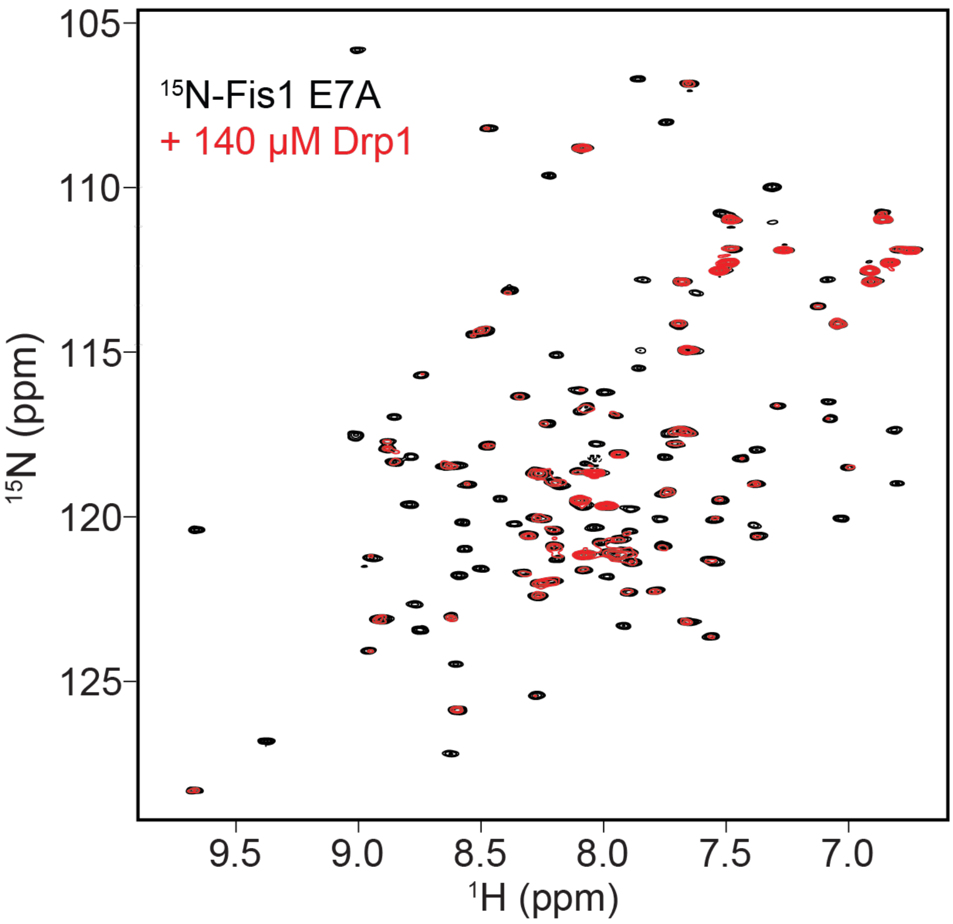

$^{15}$N-Fis1 E7A
+ 140 µM Drp1
105
110
115
120
125
$^{15}$N (ppm)
9.5
9.0
8.5
8.0
7.5
7.0
$^{1}$H (ppm)


**Figure S9.**

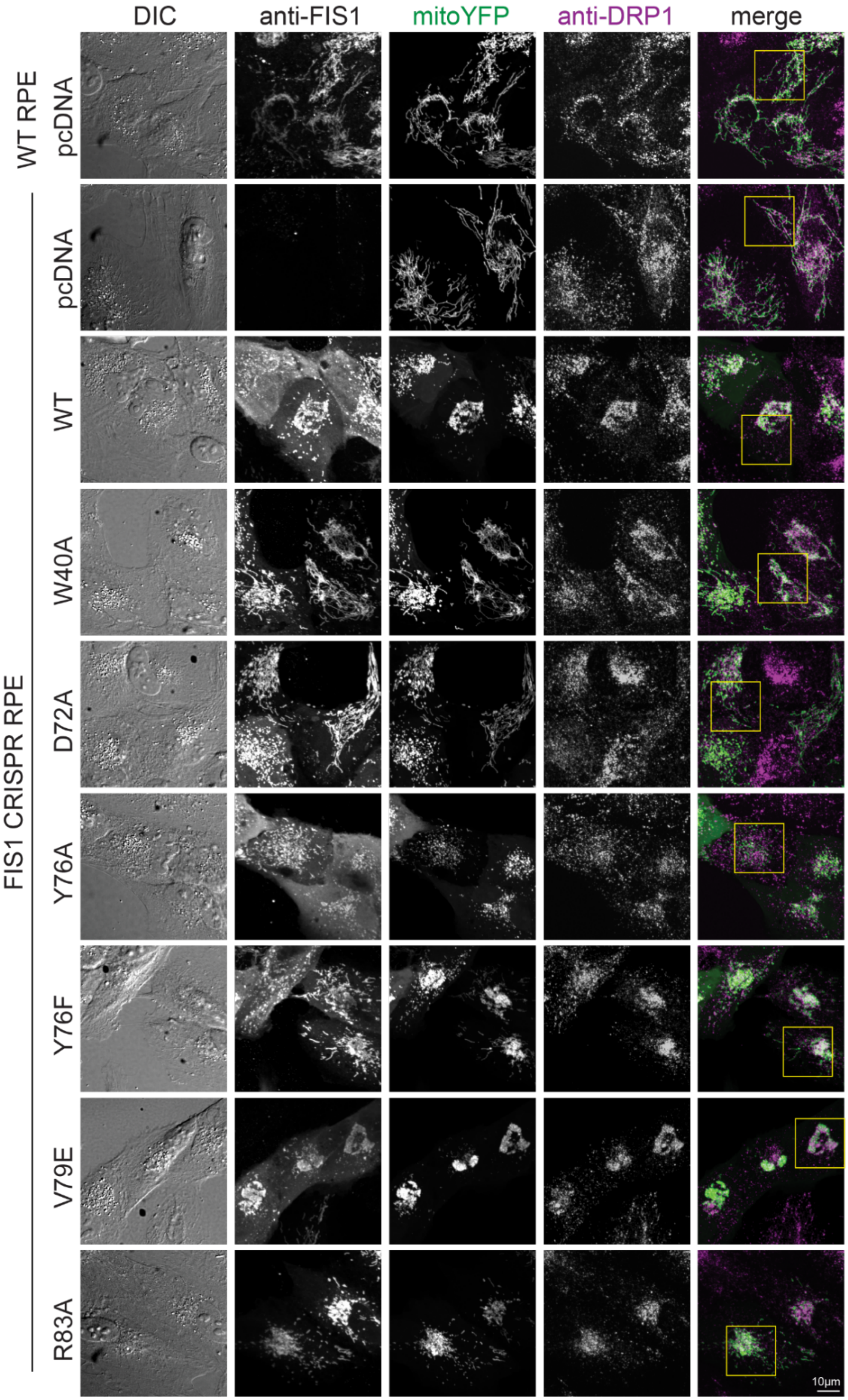
DIC
anti-FIS1
mitoYFP
anti-DRP1
merge
WT RPE
pcDNA
FIS1 CRISPR RPE
pcDNA
WT
W40A
D72A
Y76A
Y76F
V79E
R83A
10μm

**Figure S10.**

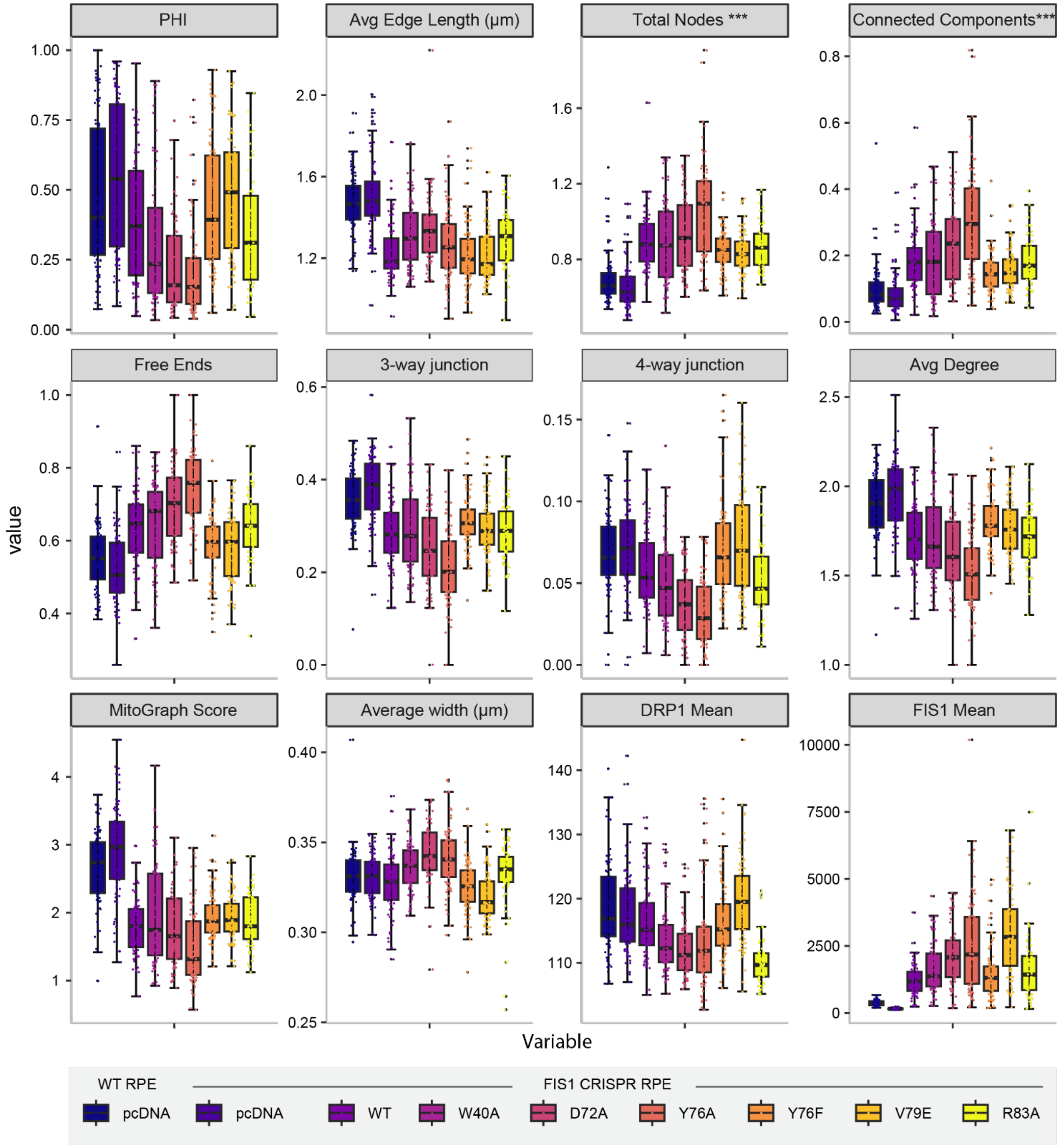
PHI
Avg Edge Length (µm)
Total Nodes ***
Connected Components***
Free Ends
3-way junction
4-way junction
Avg Degree
MitoGraph Score
Average width (µm)
DRP1 Mean
FIS1 Mean
value
Variable
WT RPE
FIS1 CRISPR RPE
pcDNA
pcDNA
WT
W40A
D72A
Y76A
Y76F
V79E
R83A

**Figure S11.**

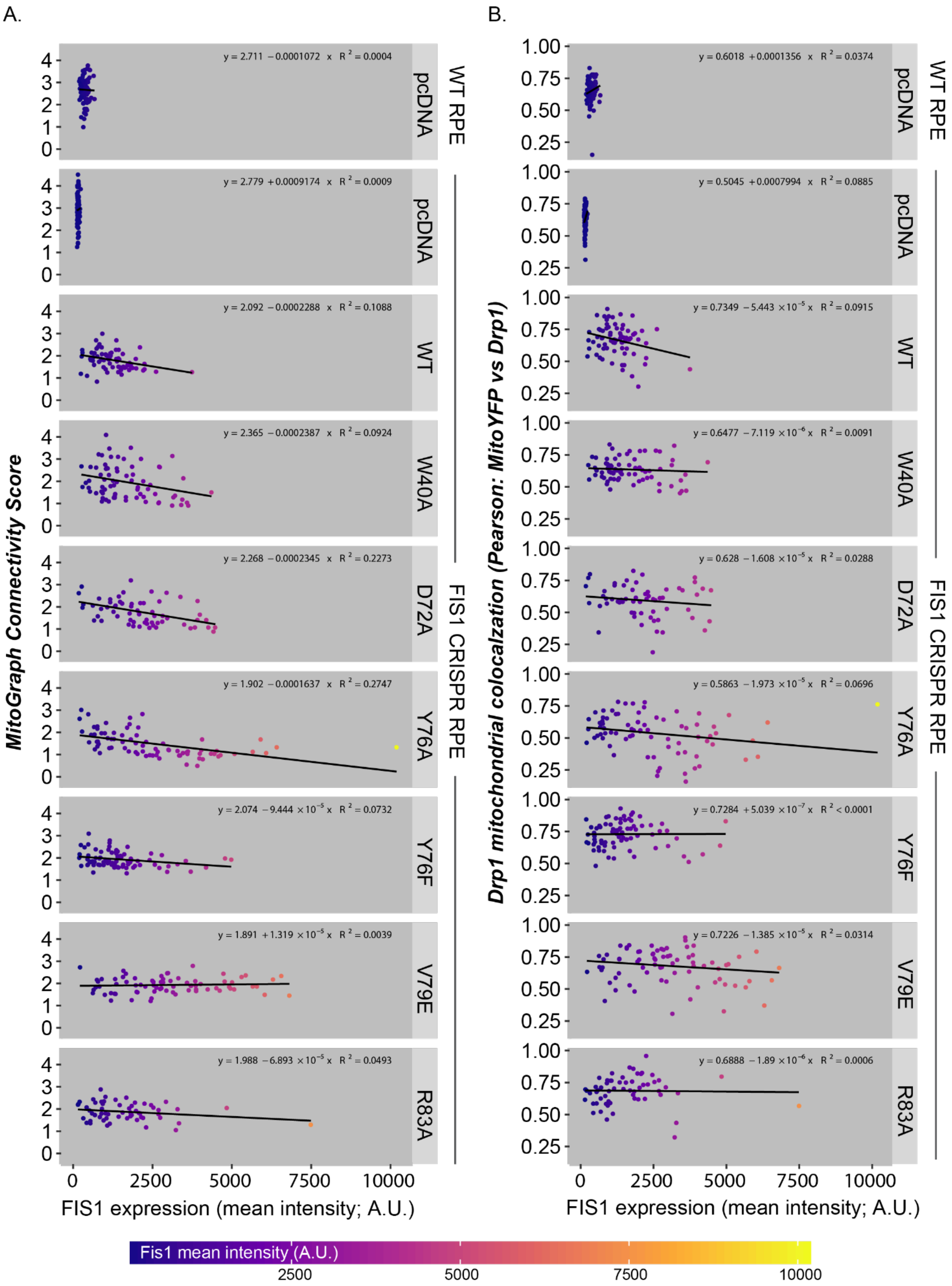

A.
B.
MitoGraph Connectivity Score
Drp1 mitochondrial colocalzation (Pearson: MitoYFP vs Drp1)
FIS1 expression (mean intensity; A.U.)
WT RPE
FIS1 CRISPR RPE
pcDNA
WT
W40A
D72A
Y76A
Y76F
V79E
R83A
y = 2.711 − 0.0001072 x R² = 0.0004
y = 2.779 + 0.0009174 x R² = 0.0009
y = 2.092 − 0.0002288 x R² = 0.1088
y = 2.365 − 0.0002387 x R² = 0.0924
y = 2.268 − 0.0002345 x R² = 0.2273
y = 1.902 − 0.0001637 x R² = 0.2747
y = 2.074 − 9.444 × 10⁻⁵ x R² = 0.0732
y = 1.891 + 1.319 × 10⁻⁵ x R² = 0.0039
y = 1.988 − 6.893 × 10⁻⁵ x R² = 0.0493
y = 0.6018 + 0.0001356 x R² = 0.0374
y = 0.5045 + 0.0007994 x R² = 0.0885
y = 0.7349 − 5.443 × 10⁻⁵ x R² = 0.0915
y = 0.6477 − 7.119 × 10⁻⁶ x R² = 0.0091
y = 0.628 − 1.608 × 10⁻⁵ x R² = 0.0288
y = 0.5863 − 1.973 × 10⁻⁵ x R² = 0.0696
y = 0.7284 + 5.039 × 10⁻⁷ x R² < 0.0001
y = 0.7226 − 1.385 × 10⁻⁵ x R² = 0.0314
y = 0.6888 − 1.89 × 10⁻⁶ x R² = 0.0006
Fis1 mean intensity (A.U.)
2500
5000
7500
10000


**Figure S12.**

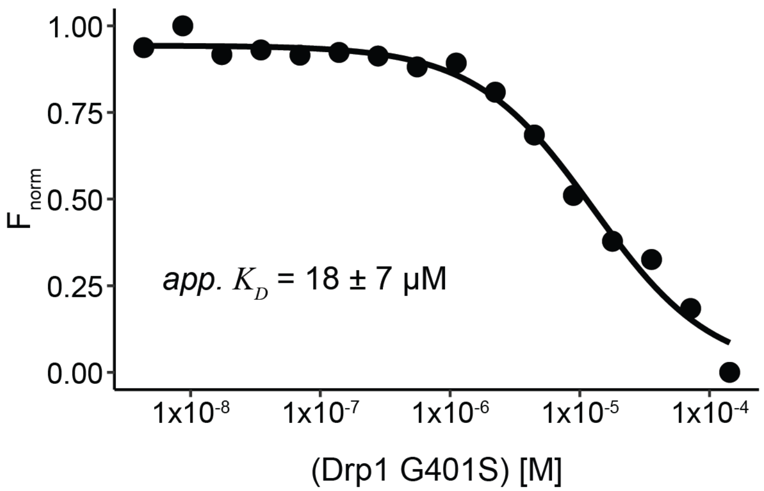
1.00
0.75
0.50
0.25
0.00
F norm
app. K D = 18 ± 7 µM
1x10-8
1x10-7
1x10-6
1x10-5
1x10-4
(Drp1 G401S) [M]

**Figure S13.**

**Table S1.** List of reagents and equipment used in this study. Source and additional notes are included.

| Item | Source | Notes |
|---|---|---|
| Chemicals | | |
| $^{15}$N ammonium chloride | Cambridge Isotopes Laboratories | |
| Avalanche-Omni | EZ Biosystems | Followed manufacturer protocols |
| DMEM-F12 | Thermo Fisher Scientific | |
| FBS | Gemini | |
| Protease Inhibitor Cocktail (EDTA free) | Roche | |
| Ni Sepharose High Performance IMAC resin | GE Healthcare/Cytiva | |
| Cy5-azide | Click Chemistry Tools | |
| BTTA | Click Chemistry Tools | |
| Cells | | |
| BL21 (DE3) pREP4 E. coli strain | Volkman lab | Ampicillin and Kanamycin resistant |
| BL21 (DE3) E. coli strain | Volkman lab | Ampicillin resistant |
| RPE (retinal pigment epithelium) | ARPE-19 (ATCC CRL-2302) | • Plated 37,500 cells/24-well<br>• Total DNA:Avalanche (1.25 µg:1 µL) |
| Antibodies | | |
| Anti-Fis1 (rabbit) | ProteinTech; 10956-1-AP (polyclonal) | Immunostaining: 1:100 (o/n @ 4˚C) |
| Anti-Drp1 (mouse) | BD; 611113 (monoclonal) | • Immunostaining: 1:100 (o/n @ 4˚C) |
| F(ab')2-Goat anti-Mouse AlexaFluor Plus 647 Cross-Adsorbed IgG (H+L) | Life Technologies A48289TR (polyclonal) | • Immunostaining: 1:500 (1 hr @ RT) |
| Goat anti-Rabbit AlexaFluor 405 Cross-Adsorbed IgG (H+L) | Life Technologies A31556 (polyclonal) | • Immunostaining: 1:500 (1 hr @ RT) |
| Plasmids | | |
| pQE30 Smt3-His$_6$-Fis1$^{1-125}$ | Hill & Volkman | • (Egner et al. 2022) |
| pQE30 Smt3-His$_6$-Fis1ΔN$^{9-125}$ | Hill & Volkman | • (Egner et al. 2022) |
| pET29b Drp1 isoform 1 TEV-His$_6$ | Hill & Volkman | • (Nolden et al. 2022) |
| pcDNA 3.1 (-) | Hill Lab | • (Egner et al. 2022) |
| pQE30 Smt3-His$_6$-Fis1$^{1-125}$ E7A | Hill Lab | • Cloned by Genscript |
| pQE30 TEV Protease S219V polyR His$_8$ | Volkman Lab | |
| pcDNA 3.1 Fis1 WT | Hill Lab | • (Egner et al. 2022) |
| pcDNA 3.1 Fis1ΔN | Hill Lab | • (Egner et al. 2022) |
| pcDNA 3.1 Fis1 E2A | Hill Lab | • Cloned by Genscript |
| pcDNA 3.1 Fis1 A3G | Hill Lab | • Cloned by Genscript |

| | | |
|---|---|---|
| pcDNA 3.1 Fis1 V4A | Hill Lab | • Cloned by Genscript |
| pcDNA 3.1 Fis1 L5A | Hill Lab | • Cloned by Genscript |
| pcDNA 3.1 Fis1 N6A | Hill Lab | • Cloned by Genscript |
| pcDNA 3.1 Fis1 E7A | Hill Lab | • Cloned by Genscript |
| pcDNA 3.1 Fis1 L8A | Hill Lab | • Cloned by Genscript |
| pcDNA 3.1 Fis1 W40A | Hill Lab | • Cloned by Genscript |
| pcDNA 3.1 Fis1 D72A | Hill Lab | |
| pcDNA 3.1 Fis1 Y76A | Hill Lab | |
| pcDNA 3.1 Fis1 Y76F | Hill Lab | |
| pcDNA 3.1 Fis1 V79E | Hill Lab | • Cloned by Genscript |
| pcDNA 3.1 Fis1 R83A | Hill Lab | |
| Hardware | | |
| Nikon Ti2 CSU W1 | Hill Lab & Hudson Lab | • Nikon Eclipse Ti2-E microscope<br>• Yokogawa confocal scanner unit (CSU-W1) (50 micron pinhole)<br>• Hamamatsu ORCA-Flash4.0 V3 sCMOS Camera<br>• 0.3 micron z-slices; 0.11 µm/pixel resolution<br>• Cells imaged using a 60 x oil objective (Plan Apo; 1.40 NA) |
| Prometheus NT.48 | NanoTemper | |
| Monolith | NanoTemper | |
| Prometheus NT.48 Series nanoDSF high sensitivity capillaries | NanoTemper | |
| Monolith Standard Capillaries | NanoTemper | |
| No. 1.5 glass bottom 24-well dishes | Cellvis | |

**Table S2.** Fis1 ConSurf scores from Fis1 NMR Structure (1PC2). A lower number indicates a higher degree of conservation.

| RES # | AA | ConSurf Score |
|---|---|---|
| **1** | MET | -1.517 |
| **2** | GLU | -0.739 |
| **3** | ALA | 0.431 |
| **4** | VAL | -0.375 |
| **5** | LEU | -0.852 |
| **6** | ASN | 1.43 |
| **7** | GLU | 1.43 |
| **8** | LEU | 1.915 |
| **9** | VAL | -0.645 |
| **10** | SER | -0.192 |
| **11** | VAL | 2.379 |
| **12** | GLU | 0.345 |
| **13** | ASP | -0.932 |
| **14** | LEU | -0.817 |
| **15** | LEU | 1.699 |
| **16** | LYS | 0.239 |
| **17** | PHE | -0.044 |
| **18** | GLU | -0.503 |
| **19** | LYS | 2.387 |
| **20** | LYS | -0.15 |
| **21** | PHE | -0.966 |
| **22** | GLN | 0.886 |
| **23** | SER | 1.019 |
| **24** | GLU | -0.81 |
| **25** | LYS | 1.538 |
| **26** | ALA | 1.293 |
| **27** | ALA | 0.835 |
| **28** | GLY | 1.38 |
| **29** | SER | 2.414 |
| **30** | VAL | 0.342 |
| **31** | SER | -0.298 |
| **32** | LYS | 0.934 |
| **33** | SER | -0.249 |
| **34** | THR | -0.687 |
| **35** | GLN | -0.541 |
| **36** | PHE | -1.303 |
| **37** | GLU | -0.633 |
| **38** | TYR | -0.874 |
| **39** | ALA | -1.31 |
| **40** | TRP | -0.542 |
| **41** | CYS | -0.784 |
| **42** | LEU | -1.589 |
| **43** | VAL | -0.963 |
| **44** | ARG | -1.04 |
| **45** | SER | -1.459 |
| **46** | LYS | 2.254 |
| **47** | TYR | 1.718 |
| **48** | ASN | 1.784 |
| **49** | ASP | 2.418 |
| **50** | ASP | -0.501 |
| **51** | ISE | -0.861 |
| **52** | ARG | 0.413 |
| **53** | LYS | -0.183 |
| **54** | GLY | -1.16 |
| **55** | ILE | -0.533 |
| **56** | VAL | 1.787 |
| **57** | LEU | -0.952 |
| **58** | LEU | -1.065 |
| **59** | GLU | -0.262 |
| **60** | GLU | 0.326 |
| **61** | LEU | -0.993 |
| **62** | LEU | -0.025 |
| **63** | NA | 0 |
| **64** | LYS | 1.397 |
| **65** | GLY | 2.399 |
| **66** | SER | 0.636 |
| **67** | NA | 0 |
| **68** | GLU | 1.461 |
| **69** | GLU | 0.071 |
| **70** | GLN | 0.174 |
| **71** | ARG | -1.222 |
| **72** | ASP | -1.17 |
| **73** | TYR | -0.632 |
| **74** | VAL | -0.876 |
| **75** | PHE | -0.887 |
| **76** | TYR | -0.933 |
| **77** | LEU | -1.111 |
| **78** | ALA | -1.141 |
| **79** | VAL | -0.521 |
| **80** | GLY | -0.857 |
| **81** | ASN | 0.01 |
| **82** | TYR | -0.394 |
| **83** | ARG | -1.169 |
| **84** | LEU | -0.569 |
| **85** | LYS | -0.074 |
| **86** | GLU | -0.051 |
| **87** | TYR | -0.92 |
| **88** | GLU | 0.891 |
| **89** | LYS | 0.926 |
| **90** | ALA | -1.458 |
| **91** | LEU | -0.978 |
| **92** | LYS | 0.372 |
| **93** | TYR | -0.266 |
| **94** | VAL | -0.134 |
| **95** | ARG | 1.238 |
| **96** | GLY | 0.673 |
| **97** | LEU | -1.099 |
| **98** | LEU | -0.795 |
| **99** | GLN | 1.949 |
| **100** | THR | 0.206 |
| **101** | GLU | -0.843 |
| **102** | NA | -0.774 |
| **103** | GLN | 1.181 |
| **104** | ASN | -1.352 |
| **105** | ASN | 1.093 |
| **106** | GLN | -1.442 |
| **107** | ALA | -1.243 |
| **108** | LYS | 1.339 |
| **109** | GLU | 0.836 |
| **110** | LEU | -1.58 |
| **111** | GLU | 0.159 |
| **112** | ARG | 2.156 |
| **113** | LEU | 0.539 |
| **114** | ILE | -0.879 |
| **115** | ASP | 0.55 |
| **116** | LYS | 0.321 |
| **117** | ALA | 0.373 |
| **118** | MET | -0.862 |
| **119** | LYS | 0.86 |
| **120** | LYS | 0.354 |
| **121** | ASP | -0.432 |
| **122** | GLY | -0.511 |
| **123** | LEU | 0.222 |
| **124** | VAL | -0.052 |
| **125** | GLY | -1.22 |
